# A quantitative rule to explain multi-spine plasticity

**DOI:** 10.1101/2022.07.04.498706

**Authors:** T. E. Chater, M. F. Eggl, Y. Goda, T. Tchumatchenko

## Abstract

Neurons receive thousands of inputs onto their dendritic arbour, where individual synapses undergo activity-dependent changes in strength. The durable forms of synaptic strength change, long-term potentiation (LTP) and long-term depression (LTD) require calcium entry through N-methyl-D-aspartate receptors (NMDARs) that triggers downstream protein signalling cascades in the dendrite. Notably, changes in postsynaptic strengths associated with LTP and LTD are correlated to changes in spine head volume, referred to as structural LTP (sLTP) and structural LTD (sLTD). Intriguingly, LTP and LTD, including sLTP and sLTD, are not necessarily restricted to the active, targeted synapses (homosynapses), and the changes in synaptic strength can spread and affect the strengths of inactive or non-stimulated synapses (heterosynapses) on the same cell. Moreover, the plasticity outcome at both homo- and heterosynapses can depend on the number of stimulated sites when eliciting multi-spine plasticity. Precisely how neurons allocate resources for implementing the changes in strength at individual synapses depending on their proximity to input activity across space and time remains an open question. In order to gain insights into the elementary processes underlying multi-spine plasticity that engages both homosynaptic and heterosynaptic changes, we have combined experimental and mathematical modelling approaches. On the one hand, we used glutamate uncaging to precisely and systematically stimulate variable numbers of homosynapses sharing the same dendritic branch whilst monitoring tens of other heterosynapses on the same dendrite. Homosynaptic potentiation of clusters of dendritic spines leads to heterosynaptic changes that are dependent on NMDAR, CaMKII and calcineurin. On the other hand, inspired by the Ca^2+^ levels hypothesis where different amounts of Ca^2+^ lead to either growth or shrinkage of spines, we have built a model based on a dual-role Ca^2+^-dependent protein that induces sLTP or sLTD. Comparing our experimental results with model predictions, we find that *(i)* both collaboration and competition among spines for protein resources are key drivers of heterosynaptic plasticity and *(ii)* the temporal and spatial distance between simultaneously stimulated spines impact the resulting spine dynamics. Moreover, our model can reconcile disparate experimental reports of sLTP and sLTD at homo- and heterosynaptic spines. Our results provide a quantitative description of the heterosynaptic footprint over minutes and hours post-stimulation across tens of microns of dendritic space. This broadens our knowledge about the operation of non-linear dendritic summation rules and how they impact spiking decisions.

## Introduction

A typical neuron in the mammalian brain carries thousands of synapses along its dendritic arbour. Spikes arriving at the synapses can drive changes in the size and molecular composition of the corresponding postsynaptic dendritic spines and induce synaptic potentiation or depression. In classical Hebbian synaptic plasticity, the change is long-lasting and thought to be specific to active synapses (homosynapses) and independent of other synapses. Interestingly, synapses that are not directly stimulated (heterosynapses) on the same postsynaptic target neuron can be modulated nonetheless to undergo synaptic strength changes of variable polarity either locally or in distant dendritic compartments by the same activity that elicit by homosynaptic plasticity (Oh et al., 2015; Engert and Bonhoeffer, 1997; Bonhoeffer et al., 1989; Fitzsimonds et al., 1997; Royer and Paré, 2003; Lynch et al., 1977; Letellier et al., 2019; Tong et al., 2021). Furthermore, the stimulated homosynaptic spines themselves are affected by the cross-talk of stimulated and unstimulated spines giving rise to complex multi-spine plasticity patterns (Makino and Malinow, 2011; Lee et al., 2016; Chirillo et al., 2019). Consequently, when more than two plasticity induction events coincide in space and time, predicting their respective magnitude is critical for spike generation that drives circuit functions (Caroni et al., 2012; Fu et al., 2012; Hensch, 2005). Various functional roles for heterosynaptic plasticity have been proposed, from homeostatic function that supports homosynaptic change (e.g. Lynch et al., 1977; Royer and Paré, 2003) to shaping the assembly of co-active inputs in many brain regions (Chicurel and Harris, 1992; Druckmann et al., 2014; Kasthuri et al., 2015; Lee et al., 2016). Despite the large body of work on heterosynaptic plasticity, the elementary processes that instruct heterosynaptic inputs and the effective spatio-temporal footprint that evolves across minutes timescale along the dendritic arbour to produce the durable changes are still not fully understood (Chater and Goda, 2021). In part, this is a combinatorial problem that needs to be studied with defined spatial and temporal distances between stimulated and unstimulated synapses across several to tens of minutes, and a problem that would benefit from a model framework to explain the observed effects.

In broad terms, heterosynaptic plasticity is promoted by homosynaptic activity that triggers a flux of Ca^2+^ ions into the postsynaptic neuron through NMDARs and voltage-gated Ca^2+^ channels (VGCCs), which then engages a wide range of Ca^2+^-dependent signalling events that spread to neighbouring spines, including the spread of Ca^2+^ itself. A mechanism involving membrane voltage signal spread, including NMDA spikes and back-propagating action potentials, can also contribute to heterosynaptic plasticity (e.g. Schiller et al., 2000; Major et al., 2008; Branco and Häusser, 2011). Secondary events, such as activity-dependent local dendritic translation, add an additional layer of regulation. Although how activity-dependent signals and new proteins are distributed amongst spines is still under debate, proposed mechanistic frameworks include the synaptic tag and capture (STC) (Redondo and Morris, 2011) and the clustered plasticity model where the influence of plasticity is confined to individual dendritic branches (Govindarajan et al., 2006).

Many candidate molecules have been identified as potential mediators of heterosynaptic signalling. These include the Ca^2+^-calmodulin dependent protein phosphatase calcineurin (Oh et al., 2015; Tong et al., 2021), the Ca^2+^-calmodulin dependent protein kinase CaMKII (Ouyang et al., 1997; Rose et al., 2009; but see Lee et al., 2009), the small GTPases Ras (Harvey and Svoboda, 2007) and RhoA (Murakoshi et al., 2011), and the diffusible gas nitric oxide (Tong et al., 2021) amongst others (reviewed in Yasuda and Murakoshi, 2011). Following homosynaptic activity, several of these molecules have been shown to diffuse away from the activated spine and spread along the dendrite segment, whereas other components such as Cdc42 and CaMKII are likely confined to the activated spine (Murakoshi et al., 2011; Lee et al., 2009; Bosch et al., 2014). Notably, CaMKII and calcineurin act in parallel to readout frequency and strength of the stimulus (Fujii et al., 2013) where CaMKII plays a role in LTP/sLTP (e.g. Silva et al., 1992; Tan and Liang, 1996) and calcineurin is required for LTD/sLTD (Oh et al., 2015; Mulkey and Malenka, 1992). Hence, protein kinases and phosphatases are likely to be critical determinants for activating and/or allocating essential plasticity components to neighbouring spines for the expression of heterosynaptic plasticity.

Overall, the current knowledge of molecular cascades highlights the existence of cooperative mechanisms that can jointly upregulate the strengths of multiple homosynapses and heterosynapses (e.g. spreading of plasticity factors and *de novo* protein synthesis) and the contribution of counter-forces that dampen the synaptic response (e.g. competition for shared protein resources). To clarify the underlying basis of inter-synaptic coordination that neurons use to integrate information received by their dendrites, we asked whether we can quantitatively predict the spatio-temporal footprint of heterosynaptic plasticity resulting from the activity of a specific set of homosynapses based on the known features of molecular players of plasticity. To this end, we combined experimental and modelling approaches and explored the minimal principle components that drive heterosynaptic spine plasticity. Using glutamate uncaging, we systematically elicited sLTP (Matsuzaki et al., 2004; Harvey and Svoboda, 2007) at variable numbers of target spines and monitored the homosynaptic and heterosynaptic spine structural dynamics over time while also testing their sensitivity to perturbing candidate heterosynaptic signalling systems. In parallel, we built a mathematical model in which the action of Ca^2+^ and protein dynamics within the dendrite results in sLTP or sLTD at homosynaptic and heterosynaptic sites depending on the context in which activity is imposed.

Our mathematical approach was motivated by a number of previously proposed models. Following the classic Bienenstock-Cooper-Monroe (BCM) sliding threshold model for LTP and LTD induction (Bienenstock et al., 1982), Lisman (Lisman, 1989) put forward an influential Ca^2+^ threshold hypothesis that was based on experimental data (reviewed in Lisman, 2001 and Evans and Blackwell, 2015), which stated that the synapses undergo LTP or LTD according to the availability of Ca^2+^. Several models have since extended the original concept. Notably, Castellani et al. (2001) and Shouval et al. (2002) considered the Michaelis–Menten kinetics of protein kinases and protein phosphatases that are directly or indirectly regulated by intracellular Ca^2+^ and promote differing LTP and LTD responses. Shouval et al. (2002) also considered the role of NMDARs as sources of Ca^2+^ for triggering LTD and LTP. However, these Ca^2+^-driven models operate on the timescale of milliseconds with an emphasis on the initial events of plasticity induction, contrary to our interest on the timescale of tens of minutes over which heterosynaptic responses develop. Therefore, we sought to generalise these models to match the timeframe of our experimental data. Moreover, using a stochastic transition approach between high and low synaptic states, Bush and Jin (2012) could predict frequency-dependent plasticity at individual stimulated spines across diverse protocols, including subthreshold depolarisation and burst pairings. Capitalising on the above models that were based on the Ca^2+^ threshold hypothesis and showed promise in explaining the different plasticity effects, we introduced a mathematical model that coupled a fast Ca^2+^-kinetics to slow protein dynamics. Interestingly, available experimental evidence pointed to the existence not only of Ca^2+^-dependent LTP and LTD regions but to a third domain, which was referred to as the “no-man’s land” (Cho et al., 2001; Nevian and Sakmann, 2006). This regime escaped existing modelling approaches because it yielded neither clear LTP nor LTD but falls between the two, depending potentially not just on Ca^2+^ but on more parameters. Taking into account the full spectrum of possible LTP and LTD outcomes and this third region, we introduced a model that avoided an abrupt LTP/LTD border and instead continuously transitioned between LTP and LTD and back. Overall, we aimed to introduce a model with a minimal number of parameters that could reproduce experimental plasticity results on the time scale of tens of minutes, along with molecular dynamics that operated also on faster time scales. To this end, we considered in our model two distinct proteins: a fast-diffusing set of Ca^2+^-binding proteins (CBP) that enact sLTP and a slower-diffusing set of proteins (*P*) activated by Ca^2+^-binding proteins that lead to an sLTD or sLTP response based on its available concentration.

Our experimental results revealed that the spine structural plasticity response was strongly affected by the number and the spatial arrangement of quasi-simultaneously stimulated spines. We observed a weaker average spine response when glutamate uncaging was triggered at more synaptic sites, whereby the size of the average spine growth was inversely proportional to the distance to its nearest stimulated neighbours. Furthermore, the spines located within a cluster of stimulated spines but were not themselves stimulated showed responses similar to their stimulated neighbours, albeit at a reduced amplitude. Crucially, the mathematical model could accurately predict homosynaptic and heterosynaptic spine dynamics as measured in our experiments. Collectively, our results provide novel insights into competitive and collaborative mechanisms that allocate plasticity factors across spines as a function of temporal and spatial distance from the sites of induction of plasticity and, in turn, determine the plasticity response at individual spines.

## Results

On the experimental side, we began our study by eliciting sLTP at target homosynapses and assessing plasticity at neighbouring heterosynaptic spines sharing a dendrite. To this end, glutamate uncaging was used to quasi-synchronously stimulate a handful of spines (*n* = 1, 3, 7, 15) in CA1 pyramidal neurons in rat organotypic hippocampal slices. Prior biolistic transfection of slices with GFP resulted in sparse labelling of pyramidal neurons that allowed visualization of the entire dendritic tree of single neurons and discrimination of individual spines of both homosynapses and heterosynapses. Quasi-simultaneous spine activation was achieved by uncaging glutamate at each spine for 4 msec and then moving to the next spine within 3 msec. This way, stimulation of a group of 7 spines, for example, was completed within 50 msec. As reported previously, robust sLTP was observed at target stimulated spines (Figure 2 and Figure S1), and the extent increase in fluorescence signal relative to the baseline signal (i.e. the amount of plasticity) at stimulated spines depended on the number of stimulated homosynapses.

To investigate the molecular mechanisms that produced a specific amount of sLTP or sLTD at heterosynapses in association with sLTP induction at homosynapses, we built a mathematical model. Our model describes the amount of plasticity relative to the baseline for each spine. At the core of our model is a combination of fast and slow mechanisms: a fast dynamics of CBP, which in turn trigger a protein redistribution that evolves from minutes to hours. Specifically, we consider experimentally reported features of Ca^2+^-dependent activation of CaMKII and the regulatory influence of calcineurin on spine plasticity. To gain an understanding of the effect of multiple simultaneous stimulation events on the amount of sLTP or sLTD induced in the homosynapses, we have constrained our model by postulating that CBP, defined by the variable *C*, consist of an initial dendritic population, *C_d_*, which is available to all spines within a certain distance and a spine-bound initial amount, *C_s,i_*, which is only available to the spine *i*. Since the stimulation process is known to increase Ca^2+^, we differentiate in our model between stimulated and unstimulated spines by assigning all stimulated spines a variable *C_s_* that is different from the corresponding variable of the non-stimulated spines. We have then tested our first model prediction against experimental data to determine if spines share resources. We quantify the total amount of CBP available to single spines upon inducing sLTP (*C_s_*) and test whether it is inversely proportional to the number of stimulation sites, *N*:

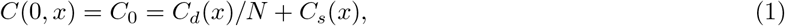

where *x* is the distance from the stimulation site, *C_0_* is the combined initial amount of *C*, and *C_d_* and *C_s_* are assumed to be maximal at *x* = 0. We note that the spatial dimension *x* is considered along the dendrite direction and is one-dimensional. After plasticity induction, CBP diffuses with a rate *a*;

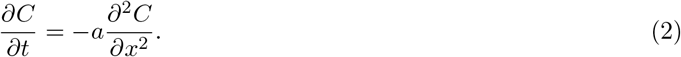

Equation (1) allows us to quantify the experimental outcome if spines are sharing resources. In other words, as sLTP is induced at an increasing number of spines, more spines will compete for factors required for potentiation such that each might undergo less potentiation. Thus glutamate uncaging at more spines reduces the average potentiation of each sLTP spine proportionally to the number of stimulated spines until a baseline is reached because as N keeps growing, the spines fall back to *C*(0) = *C_s_*. Similarly, all spines that do not receive direct stimulation (heterosynapses) have *C_s_* at their disposal such that a baseline amount of CBP-driven potentiation is assured regardless of the number of sLTP events. The diffusion equation indicates that *C* introduced at *x* = 0 diffuses to the local vicinity and leads to plasticity events at neighbouring heterosynapses if a sufficient amount of *C* is reached. Our experimental results, as well as those of others (e.g. Matsuzaki et al., 2004), indicate that significant heterosynaptic effects only arise if stimulated spines surround the heterosynaptic spines and if resource overlap from multiple nearby stimulation sites occurs.

So far, we have considered only the fast CBP proteins and their distribution across spines. Next, we introduce a slow protein resource *P* whose production is promoted by the local accumulation of CBP. Here, we follow the same approach as above and split this quantity into two terms: a dendritic pool *P_d_* that can be accessed by all spines and a spine-specific store *P_s_*. We will later treat the stimulated homosynaptic spines and heterosynaptic spines separately and assign individual *P_s_* values to these two classes. We hypothesize that the simultaneous glutamate uncaging at *N* spines will lead to *P*, which is inversely proportional to *N*:

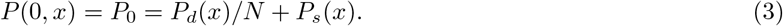

To define the time evolution of *P*, we introduce a reaction-diffusion model in which *C* drives the production of an active *P* with the rate *b_1_* and lateral diffusion *b_2_*:

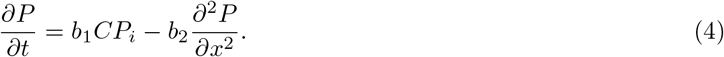

Here, the amount of *P* represents the concentration of active proteins at the corresponding spines that undergo long-term potentiation lasting tens of minutes or more. Notably, the introduction of *P* is motivated by CaMKII, which is activated by an increase in Ca^2+^ and whose activity is critical for sLTP (Miller and Kennedy, 1986; Urakubo et al., 2014). Moreover, active CaMKII can be converted into a Ca^2+^-independent, persistently active state via autophosphorylation (Glazewski et al., 2000). Experimental and theoretical evidence suggests that concentrations of active CaMKII evolve on the time scale of minutes (Fonkeu et al., 2019), which is much slower than the Ca^2+^-mediated dynamics lasting a few seconds that has been considered by previous models (Castellani et al., 2001; Shouval et al., 2002). In our model, the concentration of *P* grows as long as the accumulation rate is larger than the rate at which the protein leaves the spine. Whether a spine grows or shrinks over the long term depends on the concentrations of both CBP and *P* in line with experimental reports (e.g. Jaffe et al., 1992). If *P* is above its potentiating (i.e. LTP) peak location *ν_P_*, *P* has a net potentiating effect. On the other hand, if *P* is below the LTD peak location, *ν_D_* will trigger spine shrinkage. Early models often showed characteristically discrete states with instantaneous switching between the potentiating and depressing states. However, such a mechanism does not account for the “no-man’s land effect” (Lisman, 2001), where an intermediate region that shows neither LTP nor LTD occurs. Therefore, we include these peak locations as continuous quantities that lead to a region in the *P* concentration space where potentiation and depression cancel each other. These peak locations are then defined by a pair of Gaussians with standard unit size where the most growth and shrinkage occur at the mean. We introduce a function *F*(*P*) which ranges between +1 and −1:

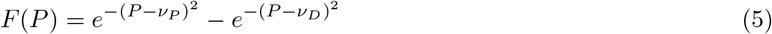

The rationale for this particular functional form of F(P) that depends only on two parameters, *ν_P_* and *ν_D_*, are as follows. We constrained the width of the Gaussians to unity, which imposes an indirect scaling on the action of protein *P*. To arrive at this, we initially considered a binary function that was positive if *P* — *ν_P_* > 0, negative if *ν_D_* <P< *ν_D_*, and zero elsewhere and that could reproduce the temporal shape of sLTP response in experiments with one, three and seven stimulated spines. However, this binary version could not be differentiated at the points *ν_D_* and *ν_P_*. Next, we increased the complexity and continuity of *F*(*P*) and fitted the following form

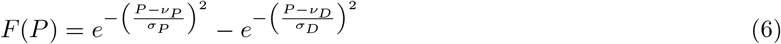

to the experimentally measured sLTP responses in plasticity induced with one, three and seven stimulated spines. Here we found that the quality of the fits (*R*^2^ values) were not significantly higher than using the functional form in Eq. 5. Therefore, we sought to continue with Eq. 5 because it offered the minimal number of parameters and mathematical continuity. Figure 1i and g illustrate how *F*(*P*) acts depending on the order of the peak locations.

**Figure 1:**
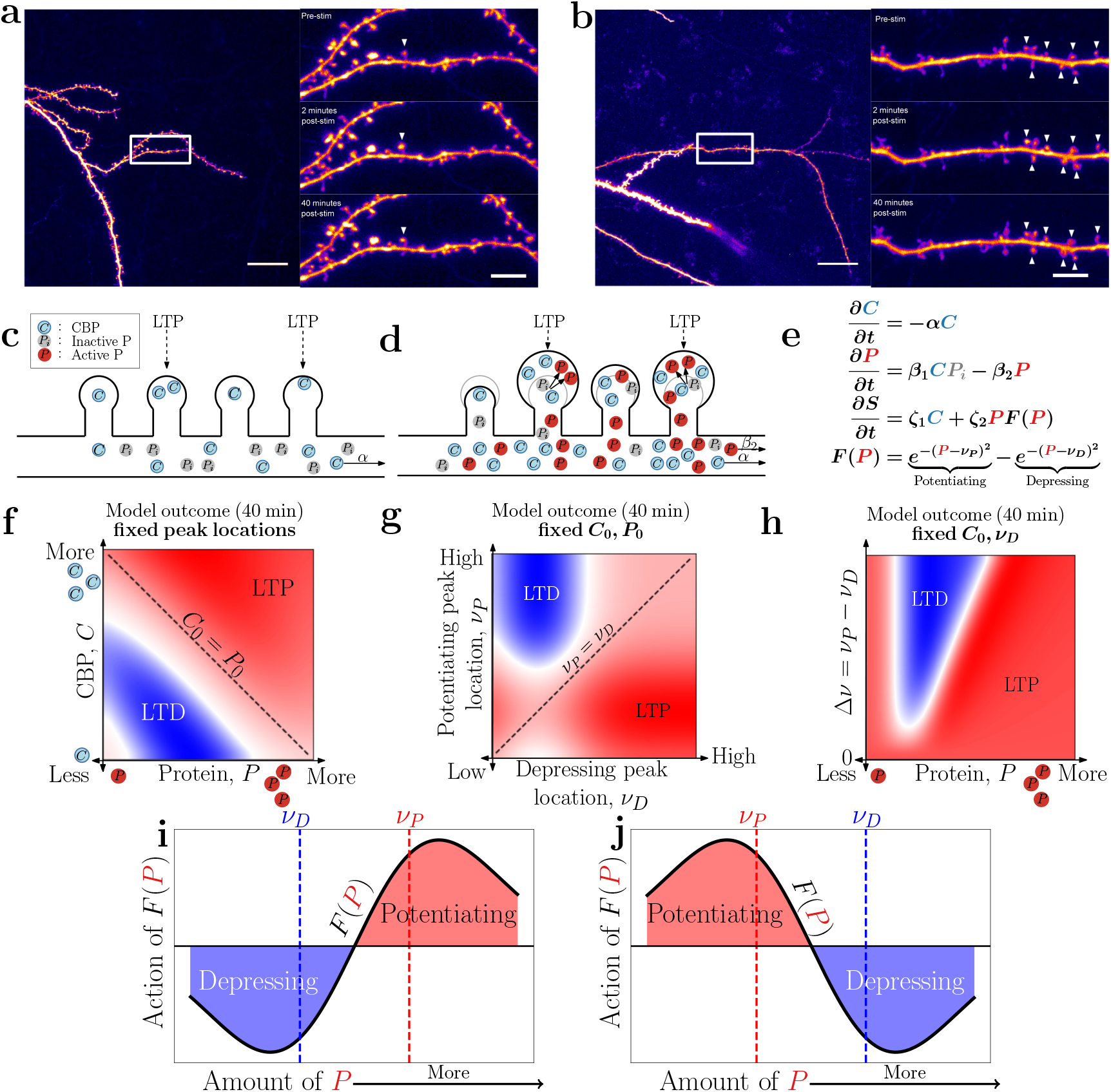
Experimental outline for examining spine dynamics along a dendrite following sLTP induction and its model description. GFP-expressing CA1 neuron used in a representative sLTP experiment targeting a single spine **a)** and seven spines, simultaneously **b)**. **Left)** Apical oblique dendrite targeted for stimulation. Scale bar = 20*μ*m. **Right)** sLTP of 1 or 7 dendritic spine(s) (white arrow heads). *Top*, before stimulation. *Middle*, 2 min post-stimulation. *Bottom*, 40 min post-stimulation. Scale bar = 5*μ*m. **c)** Before sLTP induction, a pool of CBP and inactivated *P* is available at each spine and in the dendrite. **d)** sLTP-triggered CBP increase activates the protein *P* which transitions with a rate *β_1_* from inactive *P_i_* to its active form. The activation of *P* together with CBP increase spine size. **e)** Core model equations translating CBP and *P* concentrations into spine size, color code as in (c) and (d). **f)** In the model 40 min after stimulation onset, LTP (red) occurs in a regime where either *P* or CBP are large while LTD (blue) occurs when both *P* or CBP are low. All other model parameters were kept constant, including the peak LTD/LTP locations *ν_D_* and *ν_P_*, whereas the initial concentrations *C*_0_ (*x*-axis) and *P*_0_ (*y*-axis) were varied. In **g)**, *C*_0_ and *P*_0_ where held constant while varying the potentiation and depression peak locations *ν_D_* and *ν_P_*. The dashed diagonal line refers to the 0 point, i.e. *ν*_1_ = *ν*_2_. Above the dashed line the potentiation peak location is encountered first and vice versa for the area below the line. In **h)**, *C*_0_ and depression peak location *ν_D_* are kept fixed, while *P*_0_ and the peak potentiation, *ν_P_*, are varied. The *y*-axis depicts the difference between the peak locations shown as function of the change in *P*_0_ on the *x*-axis. All three panels (f,g,h) show potentiation in red and depression in blue resulting from the model simulation at *t* = 40 min post sLTP induction. **i-j)** Action of *F*(*P*) as a function of available *P* and the LTD/LTP peak locations *ν_D_*,*ν_P_*. (left) Encountering *ν_D_* first leads initially to a depressing response before transitioning into potentiation with growing *P*. The opposite occurs when *ν_D_* > *ν_P_* (right). We note that the relative proximity ofν_D_ and *ν_P_* governs the size of the intermediate regime (previously referred to as “no-mans” land).

Next, we introduce the final component of the model, the normalized spine size *S* which is shaped by *C* and *F*:

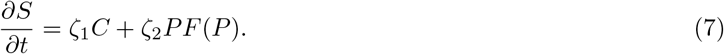

Here, *ζ*_1_ and *ζ*_2_ are parameters determining the biological susceptibility of the spines to *P*, which are held constant across all experiments regardless of the number of stimulated spines *N*. This model can be reduced to a situation where *P* is the primary driver of the synaptic plasticity by setting *ζ*_1_ = 0. However, we observed a weak sLTP in experiments with a CaMKII blocker (i.e. blockage on the actions of *P*) which motivated us to consider the *C*-mediated term that is proportional to *ζ*_1_.

Fig. 1c and d schematically summarize the model mechanisms. The spine and dendritic distribution of CBP and inactive *P* in basal conditions (Fig. 1c) changes upon the simultaneous induction of sLTP at multiple spines. CBP concentration increases in each stimulated spine along with the activation of *P* (increase in *P*: red circles in Fig. 1d).

Before continuing with the model description, here we briefly explain how moving from the spatial diffusion model to a temporal single spine model is advantageous for describing stimulated and heterosynaptic spines over time. As noted in the supplemental material, the solution to the diffusion equation at *x* = 0 (the spine location we wish to study) is equivalent (up to multiplicative constants) to the solution of the equation of a single spine with the degradation term:

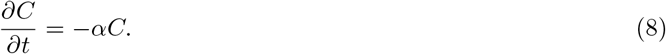

This equivalence simplifies the parameter fitting procedure, as we only need to focus on the average behaviour of a class of spines (e.g., stimulated or heterosynaptic spines) instead of considering spatial bins. The results that the single spine model produces can still be used to infer mechanisms that arise from the spatial dimension.

In the next step, we simplify the temporal equation of *C* shown in equation (8), such that *α* is now a degradation term at *x* = 0 and *P*:

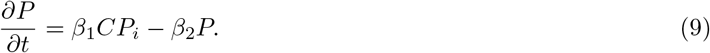

Here, *β*_1_ is the same activation term as *b*_1_ in equation (4) and *β*_2_ is a degradation term. The initial conditions (which originate from equations (1) and (3)) lose their spatial dependence and instead are simplified into a single spine model:

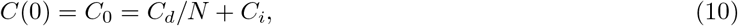

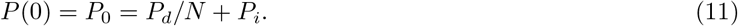

This simpler model is used for fitting the two classes of spines (stimulated and heterosynaptic) individually, and in each case, a parameter set specific to the respective class is derived. Specifically, the parameter *C*_0_ represents the quantitative amount of CBP at the stimulated spines or their spillover into heterosynaptic spines. Subsequently, whether the fit parameters of the heterosynaptic spines have the same N dependence as the corresponding fits of the stimulated spines is examined. Such a comparison should reveal whether heterosynaptic *C*_0_ and *P*_0_ both follow the 1/*N* expected from the resource competition hypothesis.

Additionally, the two model fits of the heterosynaptic and stimulated spines allow for a systematic comparison with prior literature and our own experiments. We find that when varying the different parameters, both LTP and LTD are observed: the relative states of spine sLTP or sLTD at 40 min following stimulation as a function of *C*, *P*, *ν_P_* and *ν_D_*, or the difference between *ν_P_* and *ν_D_* are shown in Fig. 1g-i (depiction of the simulated state of a spine after 40 minutes). If a spine has a low CBP value (low *C*) due to a lack of sLTP at this site and without a sufficient CBP spillover from other (nearby) activated spines, the concentration of *P* also cannot grow as a result, and the spine depresses (blue regime). This model suggests that one way to ensure robust potentiation of heterosynaptic spines is to trigger simultaneous sLTP events at multiple closely located spines such that CBP concentrations from individual sLTP events overlap at the heterosynaptic spines, thus raising *P* to reach the potentiation domain (red area: Fig. 1g). Varying the depression peak location and the relative distance between the potentiation and depression peak locations, we find robust potentiation if the difference between the two peak locations is small (Fig. 1h).

In summary, the model makes three key, experimentally testable predictions. First, raising the ratio of the LTP peak location *ν_P_* over the initially available *P*_0_ is expected to reduce potentiation at the stimulated spines on the short and long time scales. On the other hand, lowering the ratio of *ν_P_* over the *P*_0_ will increase potentiation at stimulated spines at short time scales but will leave long time scales largely unaffected because the LTP value is a non-linear function of the peak location difference *ν*_1_ — *ν*_2_. Second, our model predicts that simultaneous sLTP events at multiple closely located spines will trigger potentiation of neighbouring heterosynaptic spines due to an overlap in CBP, provided that a sufficient amount of *P* is available. Moreover, the potentiation at heterosynaptic spines grows proportionally with the number of simultaneous sLTP induction events in their immediate vicinity, particularly the long-term component of heterosynaptic potentiation. However, past a certain point when *P* becomes limiting (due to a lack of *C*), the potentiation at heterosynaptic spines will decrease proportionally to the number of sLTP events. Third, the dynamics predict that increasing the spatial distance between simultaneous sLTP events will decrease the potentiation strength until a certain distance is reached, which uncouples the sLTP events as single independent events. Finally, the model predicts that the potentiation response of stimulated spines can be inversely proportional to the number of stimulation events.

### Model summary and strategies to determine model parameters

Here, we summarize the model equations we use to fit experimental data and elaborate on our parameters-fitting procedures. The full set of model equations with their corresponding initial conditions at the time of stimulation (i.e. during baseline: defined by *t*_stim_ = 0) are listed below:

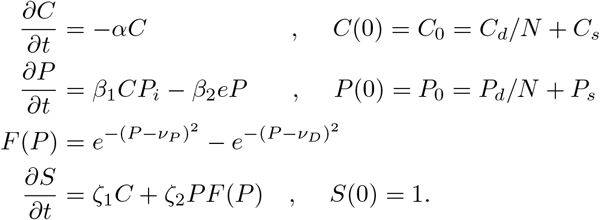

In total, our model has nine distinct parameters to be determined experimentally. These are split into two parameter classes: *(i)* cell-intrinsic parameters that are expected to remain constant across all experimental conditions and *(ii)* parameters that vary across experimental conditions, i.e., the number of stimulated sites.

The following parameters belong to the first class: *α*, *β_1_*, *β*_2_, *ζ*_1_ and *ζ*_2_ These constant parameters are fit for a representative experimental condition and then kept constant throughout all subsequent fitting involving different experimental conditions. To assess the robustness of these parameters and to confirm the presumed lack of dependency on a specific experimental paradigm, we tested their values across a different number of stimulation sites. We found good agreement of the values of these parameters across conditions as summarised in Table 1.

**Table 1:** Model parameters we assumed to be invariant across different experimental conditions. Their corresponding individual fit values are shown and validate our invariant assumption. To derive model predictions we take the average of these values. *ζ*_1_ is identically 1 for all experimental paradigms because renormalization of the governing equations eliminates this parameter.

| | Biological meaning | 1 spine | 3 spine | 7 spine | 15 spine | Mean ± 95% confidence |
| --- | --- | --- | --- | --- | --- | --- |
| $\alpha$ | Diffusion/degradation of $C$ | 0.766 | 0.854 | 0.83 | 0.788 | $0.81 \pm 0.0338$ |
| $\beta_1$ | Diffusion/degradation of $P$ | 0.927 | 0.857 | 0.904 | 0.899 | $0.896 \pm 0.0249$ |
| $\beta_2$ | Activation of $P$ by $C$ | 0.259 | 0.267 | 0.237 | 0.275 | $0.26 \pm 0.014$ |
| $\zeta_2$ | Factor of $P$ affecting $S$ | 0.712 | 0.73 | 0.648 | 0.692 | $0.695 \pm 0.0297$ |

In our model fits, four parameters, *C*_0_, *P*_0_, *ν_P_* and *ν_D_* were varied between experimental paradigms (see Table 3). Once a relationship between these parameters and the number of stimulations *N* was established, we investigated how the LTP and LTD peak positions (*ν_P_* and *ν_D_*) scaled as a function of stimulation sites. Using the obtained *C*_0_ and *P*_0_, we could also infer *C_s_*, *C_d_*, *P_s_* and *P_d_* for all experiments for the respective spine classes.

**Table 2:** Model parameters that were fit per experimental paradigm. If an entry has a *✓* then this indicates that this parameter has been altered with respect to the corresponding reference parameter set and which is noted in the “Reference set” column. A lack of a checkmark means that the parameters were carried over from the fit of the reference data set. Other fitting pathways (i.e., using the I.N.s of the three spine experiment to fit the I.N. of the seven spine paradigm) were also explored, but no discernable difference was discovered. * refers to the average fit of the control experiments across different number of stimulation events was used in this case.

| Experiment | Reference figures | Reference data set | $C_0$ | $P_0$ | $\alpha$ | $\beta_{1,2}$ | $\zeta_{1,2}$ | $\nu_{P,D}$ |
| --- | --- | --- | --- | --- | --- | --- | --- | --- |
| Single spine | Fig. 2b, c & f, Fig. 6 | N/A | ✓ | ✓ | ✓* | ✓* | ✓* | ✓ |
| Single spine (AIP) | Fig. 2c, d, g & h | Single spine |  | ✓ |  |  |  | ✓ |
| Single spine (FK506) | Fig. 2c, d, g & h | Single spine |  | ✓ |  |  |  | ✓ |
| Three spine | Fig. 3b, c & f, Fig. 6 | N/A | ✓ | ✓ | ✓* | ✓* | ✓* | ✓ |
| Three spine (I.N.) | Fig. 3b, c & f, Fig. 6 | Three spine | ✓ | ✓ |  |  |  | ✓ |
| 7 spine | Fig. 4b, c & f, Fig. 6 | N/A | ✓ | ✓ | ✓* | ✓* | ✓* | ✓ |
| 7 spine (AIP) | Fig. 4c, d, f & g | 7 spine |  | ✓ |  |  |  | ✓ |
| 7 spine (FK506) | Fig. 4c, d, f & g | 7 spine |  | ✓ |  |  |  | ✓ |
| 7 spine (I.N.) | Fig. 4b, c, e & g, Fig. 6 | 7 spine | ✓ | ✓ |  |  |  | ✓ |
| 7 spine (AIP, I.N.) | Fig. 4e & g | 7 spine (I.N.) |  | ✓ |  |  |  | ✓ |
| 7 spine (FK506, I.N.) | Fig. 4e & g | 7 spine (I.N.) |  | ✓ |  |  |  | ✓ |
| 15 spine | Fig. 3b, c & f, Fig. 6 | N/A | ✓ | ✓ | ✓* | ✓* | ✓* | ✓ |
| 15 spine (I.N.) | Fig. 5b, c & f, Fig. 6 | 15 spine | ✓ | ✓ |  |  |  | ✓ |

**Table 3:**
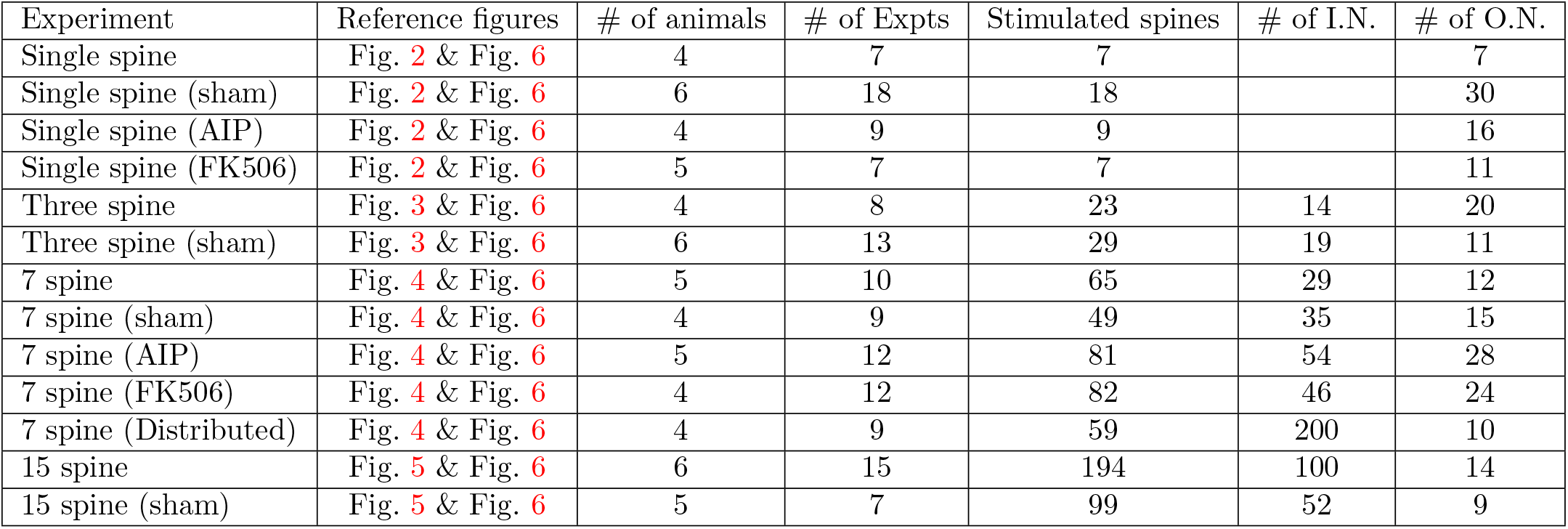
Number of studied spine types per experiment. I.N. refers to the number of inside neighbour spines not present in the single spine paradigm, and O.N. refers to the outside neighbour spines.

### Role of potentiation and depression regimes for the spine response

The predictions of our model were tested using the experimental dataset in which sLTP was induced at a varying number of stimulation spines via glutamate uncaging. The first prediction of our model states that raising the potentiation peak *ν_P_* will reduce the amount of spine growth observed at the stimulated spines on both short and long-time scales, whereas lowering *ν_P_* will increase the amount of spine growth at stimulated spines on short time scales alone. In our model, the variable *P* is motivated by Ca^2+^-dependent properties of CaMKII and its positive involvement in LTP and sLTP. To increase the sLTP peak location, we inhibited CaMKII with myristoylated autocamtide-inhibitory peptide (AIP), a specific competitive inhibitor of CaMKII (Ishida et al., 1995). Analogously, to decrease *ν_P_*, we took advantage of calcineurin that promoted sLTD and examined the consequence on sLTP triggered in the presence of calcineurin blocker FK506 (Dumont, 2000).

For visualisation, Fig. 2 shows dendritic segments containing the stimulated spine for the control and sham cases, Fig. 2 top and bottom, respectively, at 5 min before (a), 2 min after (b), and 40 min after sLTP induction (c). The stimulated spine is denoted by the white arrowhead and a dark blue ROI (see the methods section for the algorithm for the ROI assignment). The stimulated spine is denoted by the white arrowhead and a dark blue ROI (see the methods section for the algorithm for the ROI assignment). The fluorescence intensity of the stimulated spine or sham unstimulated spine was analysed before and after sLTP induction over a 55 min recording period. The mean spine fluorescence intensity change normalised to the pre-stimulation baseline (Fig. 2d) can be accurately fitted by our model to the experimentally observed sLTP, despite the presence at *t* = 30 min of a single outlier spine (see S2a). The sham non-stimulated spines remained stable over the recording period, indicating a lack of adverse effects of uncaging laser pulses. Furthermore, in the absence of MNI-glutamate, the targeted spines did not exhibit sLTP (see lower panel in Fig.2a-c).

We then repeated the experiment in the presence of AIP (5*μ*M) or FK506 (2*μ*M). Consistent with previous reports (e.g. Oh et al., 2015; Tong et al., 2021), a robust increase in early spine potentiation was observed in the presence of FK506 relative to control. In contrast, potentiation was reduced over both short and long time scales in the presence of AIP comParéd to control (Fig. 2f and h,g). The model fits (solid lines, Fig. 2f) indicate that, in the presence of FK506, there is a significant drop in both ratios, *ν_D_*/*P*_0_ and *ν_P_*/*P*_0_. In the presence of AIP, *ν_D_* increased to favour synaptic depression, which corresponded to a significantly weaker sLTP response at the stimulated spine (see Fig. 2e). These observations are consistent with our first theoretical prediction.

Next, to better understand the origin of the AIP and FK506 mediated response change, we examined the temporal dynamics of CBP, *C*(*t*) and *P*(*t*) in the model (Fig. 2g). The action of *P*(*t*) in control and the two drug conditions under AIP and FK506 helped us determine the fraction of *P* that underlies the spine plasticity. Interestingly, AIP substantially increased the depressing effect of *P*(*t*) by raising the potentiation peak location. Similarly, FK506 led to a stronger *P* response by lowering the depression and potentiation peaks, *ν_D_* and *ν_P_*, respectively (cf. Fig. 2e), and consequently leading to additional potentiation. In both cases, the dynamics of *C*(*t*) were largely unaffected by the optimisation procedure. According to the model, this implies that the blockers have little to no effect on the CBP dynamics but affect the availability of *P*.

### Differential heterosynaptic recruitment in the vicinity of multiple sLTP spines

Next, we turn to our second prediction that simultaneous sLTP events at multiple closely located spines can trigger the potentiation of neighbouring heterosynaptic spines. In the sLTP induction condition used in the present experiments, stimulation of a single spine leads to small but non-significant growth immediately after the stimulation in the heterosynaptic spines (within < 2*μ*m of the stimulated spines) that is not sustained (see Fig. S1e and f). Could the stimulation of multiple spines then be sufficiently strong to recruit heterosynaptic spines and induce longer-lasting plasticity in those non-stimulated spines? This possibility was explored using the experimental dataset in which three closely located spines were simultaneously targeted for sLTP induction (encircled in dark blue ROIs and denoted by white arrowheads in Fig. 3a - c; average distance of 4.5*μ*m). We denote all spines located between the activated spines as inside neighbour spines (I.N.) and those outside but within 2 *μ*m of the stimulated spines as outside neighbour spines (O.N.). A plot of spine fluorescence change of the stimulated spines and the immediate neighbouring I.N. spines as a function of time shows that simultaneous glutamate uncaging at three spines potentiates heterosynaptic I.N. spines at 2 min post-stimulation (Fig. 3d and 3g). Nevertheless, this heterosynaptic potentiation is not maintained over time and largely decays back to baseline (Fig. 3h). The transient potentiation of I.N. can be explained in the model by the dynamic shifts in the levels of *P* (Fig. 3e and 3f).

**Figure 2:**
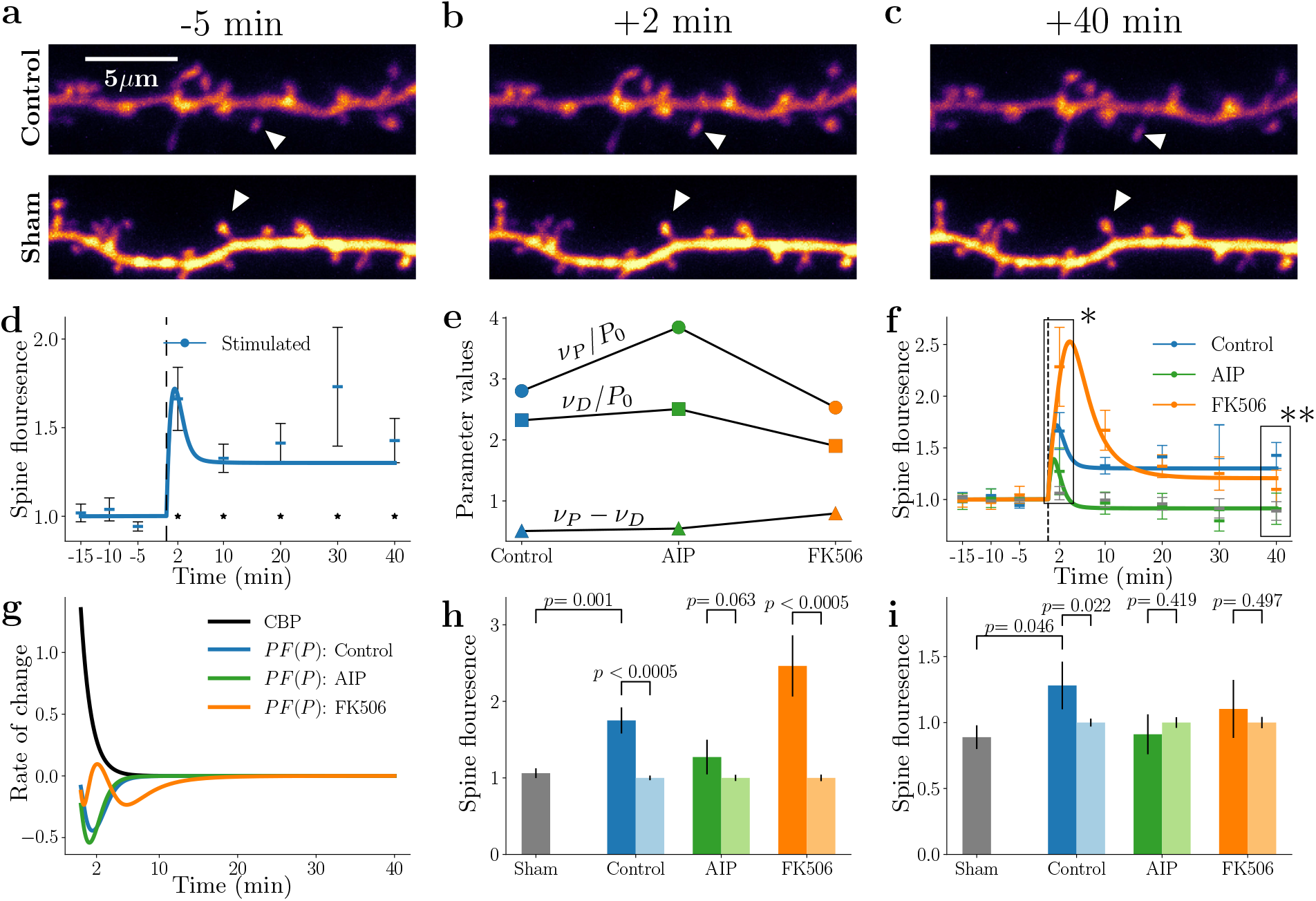
sLTP at single spines occurs via an increase in CBP. Reduction in calcineurin activity lowers the potentiation peak location, and inhibition of CaMKII raises it. **a-c)** Example images of a single spine experiment showing 5 min before, 2 min after and 40 min after applying sLTP induction protocol. The stimulated spines are marked with a white arrowhead. The top row depicts the control experiment and the bottom row the sham. **d)** Spine averaged potentiation dynamics of activated spines recorded across 40 min, averaged across sLTP experiments and normalised to pre-sLTP baseline. Superimposed is a model fit (blue line), * denotes *p* < 0.05 significance relative to sham. **e)** Parameter fits of the peak LTP/LTD locations and initial amounts of *P* for each of the experimental conditions. Calcineurin and CaMKII and their blockers, AIP and FK506, respectively, control potentiation and depression peak location and the amount of activated protein *P*. **f)** Temporal dynamics of sLTP in control conditions and in the presence of the CaMKII and calcineurin blockers. The data in the boxes marked with asterisks (*, **) are displayed for detailed comparison in figures *g* and *h*, respectively. **g)** Temporal evolution of model parameter CBP and *PF*(*P*) derived from fits in (e) across control, AIP and FK506 conditions. **h)** Normalised growth of all experimental conditions at *t* = 2 min post-sLTP (lighter bars represent the baseline before stimulation). **i)** Normalised growth of all experimental conditions at *t* = 40 min (lighter bars represent the baseline before stimulation).

**Figure 3:**
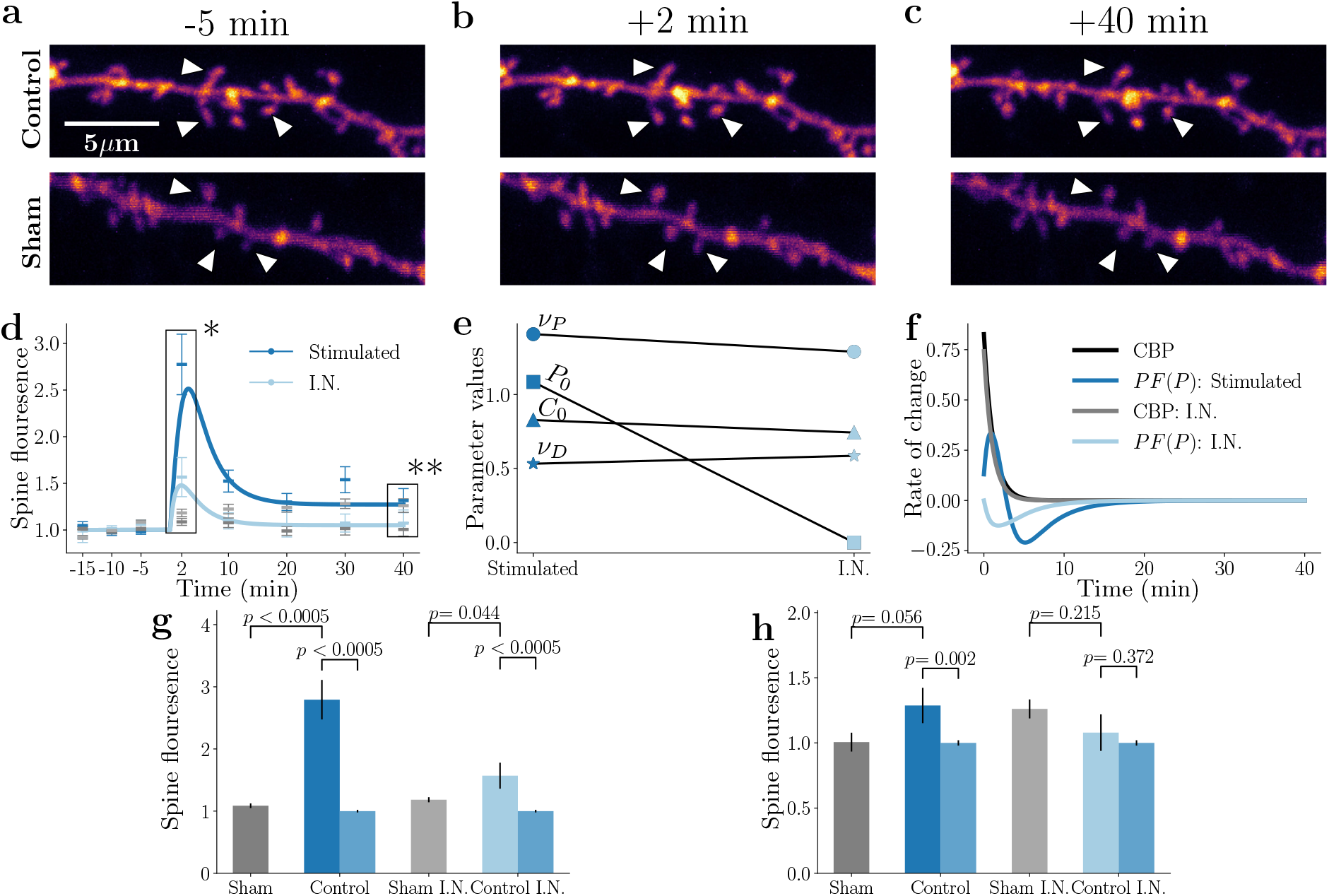
sLTP in three spines is sufficient to co-potentiate immediate neighbours (I.N.), and the amount of potentiation can be explained by a spill-over of CBP from stimulated spines. **a-c)** Dendritic anatomy of the three spine experiment for 5 minutes before, 2 minutes after and 40 minutes after a sLTP-induction. The stimulated spines are denoted by a white arrow. The top row shows the control experiment and the bottom row the sham. **d)** Normalised synaptic changes observed in the stimulated spines alongside changes in the neighbour spines and model fits. The data in the boxes marked with asterisks (*, **) are displayed for detailed comparison in figures *g* and *h*, respectively. **e)** The difference in fitted model parameters for the stimulated spines (stimulated) and immediate neighbours (I.N.). **f)** Dynamics of *C* and *F*(*P*) given by the fitted model for the stimulated and inside neighbour spines. **g)** Normalized spine growth across experimental conditions at *t* = 2 minutes, immediately after simulation. * denotes *p* < 0.05. The medium blue color for the control and control I.N. refers to the baseline values pre-stimulation of those spines. **h)** normalised growth of all experimental conditions at *t* = 40 minutes.

We next sought to test the impact of further increasing the number of spines stimulated for sLTP induction on the plasticity of stimulated and I.N. spines and analysing the experimental data with model fits. Upon stimulating seven spines, at 2 min post-stimulation, potentiation of heterosynaptic I.N. spines was still observed, whereas potentiation of targeted spines appeared slightly attenuated (Fig. 4d). Comparing the heterosynaptic I.N. response between three and seven homosynaptic spine stimulation (cf. light blue lines in Fig. 3d versus Fig. 4d), we find a decreased level of potentiation in the heterosynaptic I.N. spines that are surrounded by seven rather than three stimulated spines. This indicates that heterosynaptic potentiation, at least for those spines within the stimulated spine cluster, is reduced as the number of simultaneously stimulated spines increases. Keeping this seven-spine stimulation condition, we next studied the effects of AIP and FK506 on heterosynaptic I.N. spine plasticity similar to the single-spine stimulation case.

**Figure 4:**
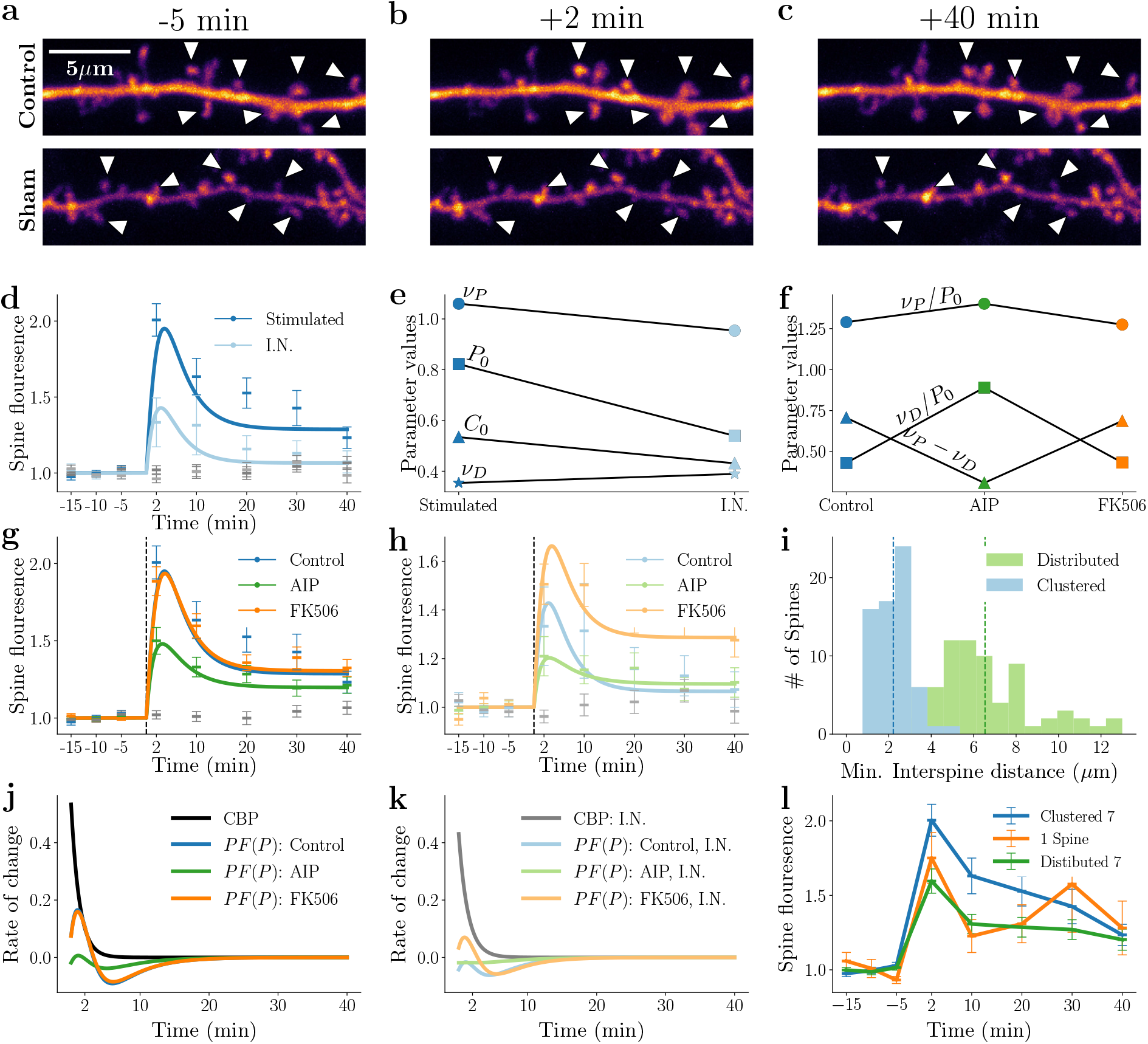
Growing distance between spines and competition for resources leads to a lower sLTP response when seven spines are stimulated simultaneously. **a)-c)** Dendritic anatomy of the seven spine experiment for 5 min before, 2 min after and 40 min after inducing sLTP. The stimulated spines are marked with a white arrow head. The top row depicts the control experiment and the bottom row the sham. **d)** Normalised spine changes in stimulated spines (dark blue) and inside neighbour (I.N.) spines (light blue), model fits are superimposed. **e)** Model parameters for stimulated and inside neighbour spines. **f)** Changes in model parameters in control and in the presence of AIP and FK506. **g)** Temporal dynamics of stimulated spines across drug conditions, model fits are superimposed. **h)** Temporal dynamics of I.N. spines in control, AIP and FK506; model fits are superimposed. **i)** Distribution of the minimum inter-spine distance between stimulated spines, a quantity related to the size of the total cluster. **j)** Model dynamics of CBP and *F*(*P*) of the stimulated spines for the control, AIP and FK506 conditions. **k)** Model dynamics of CBP and *F*(*P*) of the inside neighbour spines for the control, AIP and FK506 conditions. **l)** Normalized spine changes of stimulated spines for the clustered seven spines (blue), distributed seven spines (green) and single spine (orange) experiments.

First, for heterosynaptic I.N. spines, the level of potentiation at 2 min post-stimulation was increased by FK506 and reduced by AIP (Fig. 4g). Studying the changes in the best-fit model parameters across the heterosynaptic I.N. spine responses relative to those of stimulated spines (Fig. 4e), the level of *P* was lower at the heterosynaptic spines relative to their stimulated neighbours. The reduced *P* leads to a higher *ν_D_* /*P* ratio that dampens the overall synaptic response because the synaptic dynamics are further away from the potentiation domain around *ν_P_*. Moreover, a slight increase in the location of the depression peak *ν_D_* (Fig. 4e) further contributes to the dampening of potentiation. We also note that the CBP level *C*_0_ is lowered in the heterosynaptic I.N. spines relative to the stimulated spines. Such a situation could be potentially explained by consideration of the spatial distance between the stimulated spines, that is, the Ca^2+^ source and the heterosynaptic spines where some diffusional loss of CBP is expected. A lower CBP level, in turn, decreases the level of *P*_0_, which results in a smaller potentiation amplitude at heterosynaptic spines.

Overall, both AIP and FK506 showed lasting effects on the stimulated and heterosynaptic spines. AIP decreased the short and long-term potentiation for both the stimulated and heterosynaptic I.N. spines. This effect could be explained in our model by a change in ratio (*ν*_1_ — *ν*_2_)/*P*_0_ (Fig. 4f). FK506 led to a higher extent of short and long-term potentiation for stimulated and heterosynapses. Interestingly, in the presence of FK506, no further increase in the average spine growth of the stimulated spines was observed (Fig. 4g). In contrast, the heterosynaptic spines showed a larger degree of potentiation (Fig. 4). This can be explained in our model by the limited availability of *P* at the directly stimulated spines due to the increase in the number of stimulated spines, and even upon suppression of calcineurin activity that counteracts potentiation, stimulated spines cannot recruit additional *P* to further potentiate inputs, which is otherwise possible when only one or three spines are stimulated (see Fig. 2 and Fig. 3). In contrast, the heterosynaptic I.N. spines are facilitated to undergo potentiation by recruiting a small amount of additional *P* when relieved of activity that counteracts potentiation by FK506.

Next, comparing single, three and seven spine stimulation paradigms, we sought to test our model prediction that if the distance between the stimulated spines is kept small, then an increase in the number of stimulated spines should result in a larger potentiation due to accumulation of CBP and *P*. Consistent with the model, simultaneous potentiation of seven spines with an average inter-spine distance of 2 *μ*m (“clustered spines”; e.g. Fig. 4a) is larger than that of individually stimulated spines (Fig. 4l: see also Fig. 2d and Fig. 4d). This potentiation advantage disappears if the stimulated spines are spaced further apart with an average inter-spine distance of 6 *μ*m (“distributed spines”). The effect of spatial confinement of simultaneously stimulated spines is summarised in Fig. 4l, where the potentiation for seven distributed spines is not significantly different from the single spine stimulation. Thus, more simultaneous and clustered sLTP events increase the size of early potentiation, which is consistent with the prediction, but only if they occur close in space and time. Our model provides a mechanistic framework for the spatial limit of simultaneous sLTP induction: more simultaneous events within a confined area release more *P* due to the overlapping Ca^2+^ events and, in turn, promote a larger initial sLTP size (see Fig. 4j and k). However, if the inter-spine distance grows, the spines will decouple because the spread of CBP and *P* from individual sLTP spines do not travel fast enough to influence each other or are attenuated before they can interact.

Furthermore, in line with the model prediction, a comparison of the spine response following simultaneous stimulation of seven versus three spines (Fig. 3d) indicates that simultaneously triggering a larger number of clustered sLTP events leads to a smaller response at the stimulated spines. Specifically, the level of potentiation of both the early and late phases is reduced when seven instead of three spines are stimulated. Such a result implies that the stimulated spines have passed a certain threshold where CBP and *P* are no longer available to support additional growth. In other words, the local competition for resources could lower the degree of potentiation spines undergo following synaptic activity.

### Simultaneous glutamate uncaging of fifteen spines reveals strong competitive and cooperative components at play at the stimulated and heterosynaptic spines

Thus far, our model can reasonably explain the experimental results. Upon considering the cases for simultaneous stimulation of one, three and seven target spines, we observed that the fast potentiation component was the highest when three spines were simultaneously stimulated, and this component declined in the case of seven spines (cf. Fig. 3, Fig. 4 and Fig. 6). In order to assess the robustness of the model and gain further insights into the dynamics of model parameters, we sought to push the envelope further and simultaneously potentiated fifteen closely located spines. We then monitored spine plasticity at stimulated spines and their heterosynaptic neighbours. In particular, we wondered if the fast potentiation component declined further when fifteen spines were stimulated simultaneously and whether the parameters *P*_0_ and *C*_0_ at the stimulated and heterosynaptic spines could reveal a systematic dependence of initial *C*_0_ on the number of stimulated spines.

**Figure 5:**
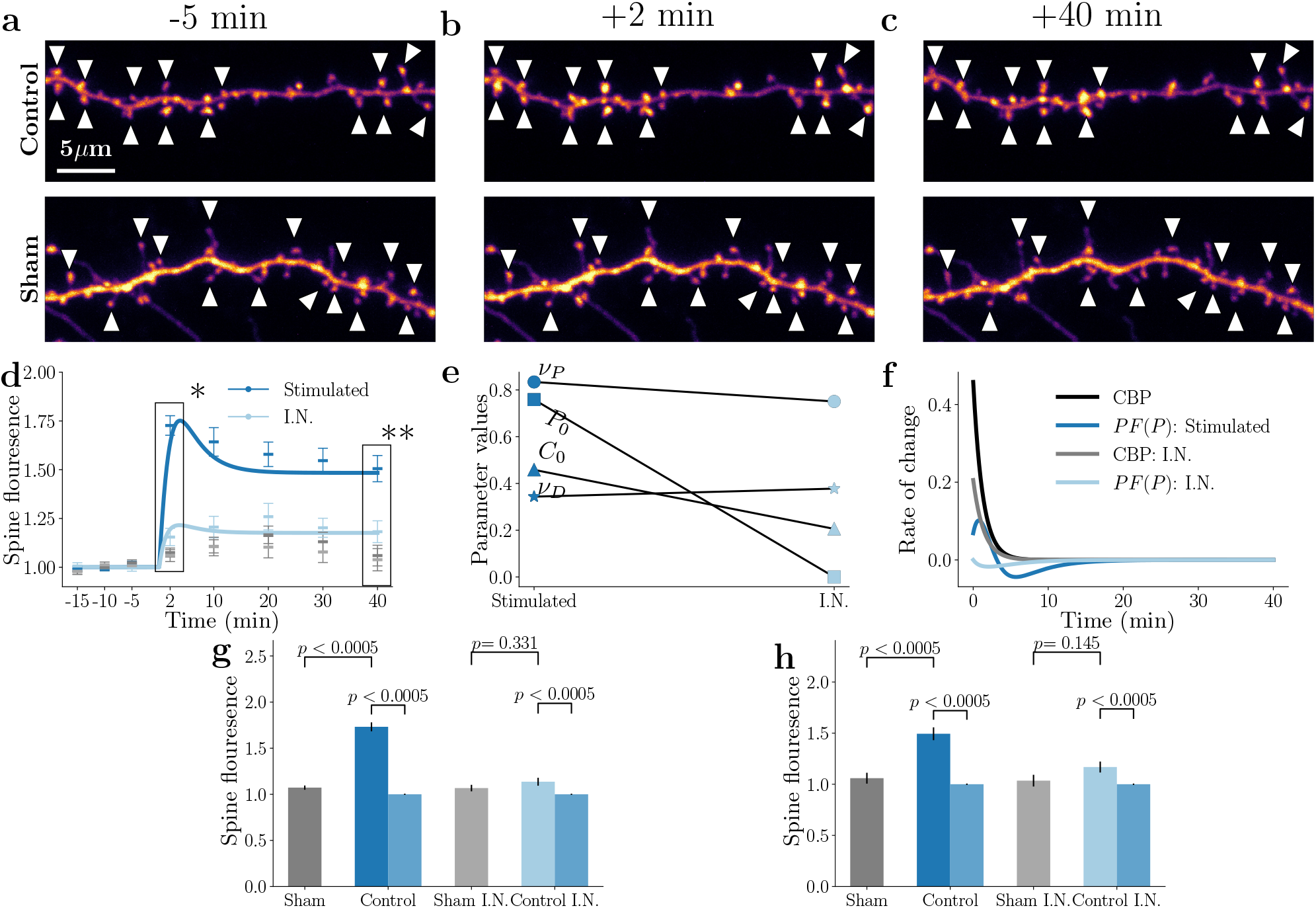
Simultaneous induction of sLTP at fifteen spines reduces the amount of spine growth in the short time but increases spine strength for long time scales. **a)-c)** Dendritic anatomy of the fifteen spine experiment at 5 min before, 2 min after and 40 min after inducing sLTP. The stimulated spines are marked with a white arrow head. The top row depicts the control experiment and the bottom row the sham. **d)** Normalised synaptic changes observed in the stimulated spines alongside changes in the neighbour spines and model fits. The data in the boxes marked with asterisks (*, **) are displayed for detailed comparison in *g* and *h*, respectively. **e)** The difference in model parameters for the stimulated spines (stimulated) and immediate neighbours (I.N.). **f)** Dynamics of *C* and *F*(*P*) given by the fitted model for the stimulated and I.N. spines. Insets represent the cumulative dynamics. **g)** Normalized spine growth across experimental conditions at *t* = 2 min, immediately after simulation. * refers to *p* < 0.05. **h)** normalised growth of all experimental conditions at *t* = 40 min.

**Figure 6:**
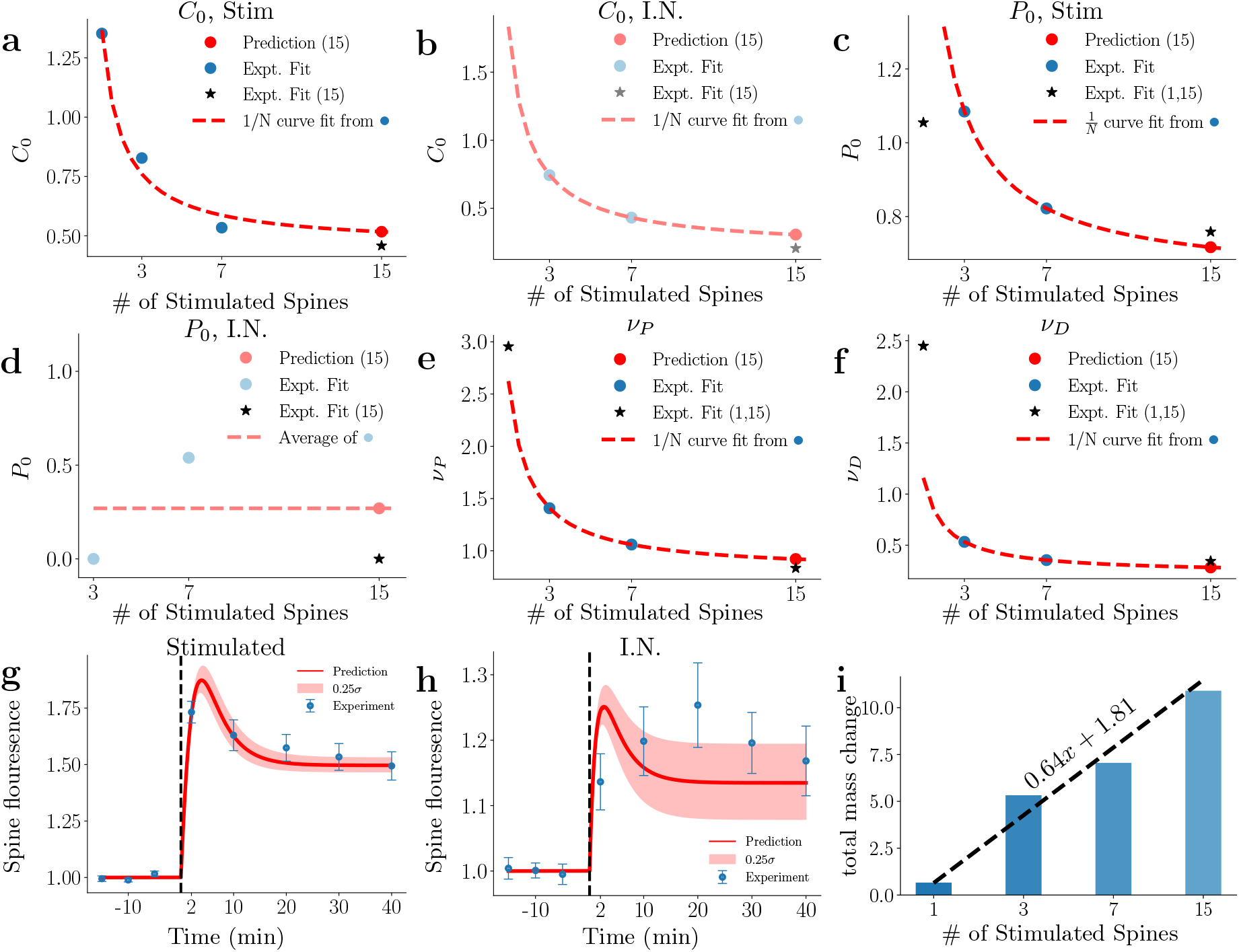
Our model predicts that a mix of spine competition and cooperation define the dynamics across a varying number of simultaneous sLTP events. **a)-b)** Decline in the initially available amount of CBP, *C*_0_, in the stimulated (a) and inside neighbour spines (b) is inversely proportional to the number of induced sLTP events (1/*N* stimulated spines). We used this 1/*N* scaling to predict (from the 1,3 and 7 spine experiments) the amount available to the stimulated spines in the 15 spine experiment (denoted by •) and which shows good agreement with the fitted parameters (⋆). **c)** Decline in the initially available amount of P, *P*_0_, in the stimulated spines is inversely proportional to the number of induced sLTP events. The prediction from the values of the 3 and 7 spine experiments were in good agreement with the directly fitted value of the 15 spine experiment. **d)** Fitted model parameters for the initially available amount of P, *P*_0_, for the inside neighbour spines. **e)-f)** Decline in the peak potentiation location, *ν_P_* (e), and peak depression location, *ν_D_* (f), is inversely proportional to the number of induced sLTP events (1/*N* stimulated spines). This effect can be used to predict (from the 3 and 7 spine experiments) the peak locations of the spines in the 15 spine experiment (denoted by o) and which in good agreement with the fitted parameters (*). **g)-h)** Model prediction for the stimulated (g) and inside neighbour (h) spines for the experiment with 15 sLTP sites. **i)** Total synaptic growth (integrated across all stimulated spines of an experiment relative to baseline) vs # stimulated spines demonstrates a cooperative increase in synaptic strength even though sLTP amplitude at the individual sLTP spines declines as a function of # stimulated spines, see Fig. S1.

Fig. 5a-c illustrates a representative example of the layout of the fifteen spine experiments. The fifteen stimulated spines (dark blue ROIs with white arrowheads) are located on a single dendritic branch and interspersed amongst the unstimulated heterosynaptic I.N. spines (encircled by light blue ROIs). The normalised spine fluorescence changes at the stimulated and heterosynaptic I.N. spines across time are shown in Fig. 5d. ComParéd to three and seven spine stimulation experiments, in both the stimulated and heterosynaptic I.N. spines, the level of potentiation at 2 min post-stimulation is reduced, whereas, at 40 min, the level of potentiation is maintained at a higher level (Fig. 5g and h). A plot of the model parameter values (Fig. 5e) illustrates that the initial *C*_0_ is similar in the stimulated and the heterosynaptic I.N. spines. However, the values of *C* and *F*(*P*) are approximately half of the values observed in the experiments with three spines (cf. Fig. 3e). Interestingly, *C*_0_ declines steadily from the three, seven, and fifteen spine experiments for the stimulated and heterosynaptic I.N. spines. From the three to the seven spine experiments, *C*_0_ decreases by 36.4% and 42% for the stimulated and I.N. spines, respectively; from the seven to the fifteen spine experiments, *C*_0_ decreases to 14.3% and 52.3% for the stimulated and I.N. spines, respectively. We will later take advantage of this drop to test for spine competition as described by equations (1) and (3).

The dynamics of *P* across spines show that the *P*_0_ immediately after the start of the glutamate uncaging is elevated in the stimulated spines. However, the heterosynaptic spines do not profit from this increase, and their *P*_0_ value is close to zero. This suggests that the heterosynaptic I.N. potentiation dynamics are initiated by the CBP spillover from stimulated spines, which raises the local concentration of *P* to further potentiate the heterosynaptic I.N. spines. We interpret that a time delay for CBP-dependent accumulation of *P* to a level sufficient to support robust sLTP at I.N. spines contributes to the observed reduction of the early potentiation component at 2 min post-stimulation. However, once a stable level of *P* is achieved, *P*-driven sLTP that is predominant by 40 min is comparable in size to that observed for the stimulation of seven spines (see Fig. 4).

Overall, these findings show that the number of spines simultaneously stimulated to undergo sLTP is a potent predictor of heterosynaptic potentiation at spines that share the dendritic locale: the more stimulated spines undergo sLTP, the more pronounced sLTP is also observed for heterosynaptic I.N. spines. Additionally, our model predicts that spine competition depletes the local stores of *P* and Ca^2+^ needed for spine potentiation and will therefore reduce the fast potentiation response at the simultaneously stimulated spines proportionally to the number of stimulated spines.

### Comparing hetero- and homosynaptic potentiation across varying number of glutamate uncaging sites

Next, we summarise the spine plasticity dynamics across all our experiments. We simultaneously stimulated a variable number of spines on a local dendritic branch and addressed the similarities and differences to help clarify the nature of underlying processes captured in our model. We have systematically comParéd *C*_0_, *P*_0_, and potentiation and depression peaks *ν_P_* and *ν_D_*, respectively, across one, three, seven or fifteen glutamate uncaging events. By comparing the plasticity outcomes across different number of spine stimulation sites, we can obtain insight into how, on the one hand, the initial quantities of *C* and *P* vary and, on the other hand, we can observe how the assumed competition of resources affects the location of depression and potentiation peaks in our model.

First, we studied the dependence of the initial values of *C*_0_ and *P*_0_ as a function of the number of stimulation sites, using equations (1) and (3). Our first step was to find values of the synaptic stores *C_s_*, *P_s_* and dendritic *C_d_* and *P_d_* that are constant across the paradigms involving different number of stimulated spines. We considered *C* in the stimulated spines, shown in Fig. 6a. Here, the blue circles represent the fitted parameters for the 1, 3 and 7 spine experiments, while the red circles denote the extrapolated parameters for the 15 spine experiment, which have been obtained by using the 1/*N* fit of the 1, 3 and 7 spine experiments (red dashed line). The simulated dynamics of the 15 spines agreed well with the values obtained from fitting this 15 spine experiment directly (black star). From this curve, we found the values of *C_s_* and *C_d_* to be independent of the number of stimulation sites. Furthermore, an equally good agreement was found when considering Fig. 6b. This supports the utility of approximating the spatial model via a simpler single-spine model. Moreover, the reasonable fit further underscored that the *C* initially available to the I.N. spines was due to the overlap from other stimulated spines. If the opposite were true, the 1/*N* would not be a fit due to a significantly larger number of heterosynaptic spines competing for *C*. Fig. 6c shows *P*_0_ corresponding to the stimulated spines. Here, we see a similarity to the behaviour of *C*_0_, except that the single spine paradigm is below the 1/*N* fit. These fits suggest that our candidate molecule *P* requires multiple stimulation events for *C* to become active and could be subsequently redistributed among the spines. Figure 2g shows the temporal activation of *P* and indicates that the action of *P* contributed a small depressing component while the spines experienced net potentiation. For heterosynaptic I.N. spines, we observed that the dynamics of *P* was dominated by the spillover of *C*_0_ (Fig. 6d). Since the initial value, *P*_0_, at the heterosynaptic I.N. spines is small and fluctuates across experiments; it may be noisy. At the same time, the level of *C*_0_ was comparable in size to that of the stimulated spines, suggesting that *C*_0_ may be the dominant source of potentiation for the heterosynaptic I.N. spines.

Turning our attention to *ν_P_* and *ν_D_*, we first kept these quantities constant across the different stimulation paradigms. However, this restriction could not capture the observed short-time plasticity dynamics. Thus, we allowed these quantities to vary as the number of stimulation sites *N* changed. We observed a decline in *ν_P_* and *ν_D_* that was inversely proportional to the number of stimulated spines (Fig. 6e,f). This suggests that the spines lowered their potentiation and depression peak locations to offset the increasingly limited resources and to promote potentiation. Let us note that we used the same potentiation and depressing domains (same peak locations *ν_P_* and *ν_D_*) for both stimulated and heterosynaptic spines for consistency reasons and to allow detailed parameter comparison across stimulated and heterosynaptic spines. We hypothesized that both *ν_P_* and *ν_D_* had a component that stayed constant as the number of stimulation sites varied

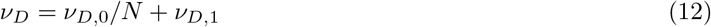

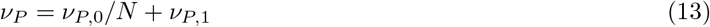

Here, *ν*_*P*/*D*,0_ represents the variable component that rises or falls in response to the amount of available *P* while *ν*_*P*/*D*,1_ is constant. Fitting *ν_P_* and *ν_D_* for the experiments with 1, 3 and 7 stimulated sites, we were able to predict the dynamics for the experiment with 15 stimulation sites for both stimulated and heterosynaptic spines (Figures 6g,h) accurately. We note that the uncertainty of the dependence on *P*_0_ for the cluster spines makes the prediction less reliable. Using the *ν*_*D*/*P*,0_ and *ν*_*D*/*P*,1_ functions we obtained from our fits we observed that these quantities are robust across stimulated and inside neighbour spines (c.f. Fig. 3e, 4e, 5e). This suggests that the only variables that change between homo- and heterosynapses (at least I.N.) are the initially available *C*_0_ and *P*_0_.

Finally, we note that the above 1/*N* dependence on the resources supports the competition for resources across spines. However, do the available resources stay constant across experiments with varying uncaging sites, or do spines cooperate to generate more resources that are then shared among all potentiating spines? To address this question, we have studied the sum total of the changes in the size of spines along the dendritic locale of stimulated spines within the field of view, including those of the stimulated spines themselves. We find that the total area increases linearly with the number of stimulated spines (Fig. 6i). If spines were only competing for a finite and constant resource, we would have expected an approximately constant integrated spine area. The linear growth proportional to the number of uncaging events could suggest the recruitment of additional resources, for example, a collaborative activation of additional CBP stores that allows the spines to potentiate.

## Discussion

Previous work exploring heterosynaptic dynamics on single dendrites has demonstrated a mixture of compensatory and non-compensatory forms of plasticity, dependent on the stimulation paradigm, age of preparation, and cell type (e.g. Engert and Bonhoeffer, 1997; Tong et al., 2021; see introduction) and has typically employed a fixed stimulation protocol to induce homosynaptic plasticity.

Here we systematically varied the number of targeted inputs from 1 to 15 spines and both directly imaged and modelled the impact of triggering sLTP on homosynaptic and heterosynaptic spine plasticity. We found that simultaneous homosynaptic potentiation of clusters of inputs led to the spreading of plasticity to nearby spines. Upon stimulating a higher number of homosynaptic inputs, compensatory mechanisms began to limit the magnitude of potentiation. Additionally, we identified a set of parameters in the context of our model that enabled us to predict plasticity behaviour at different experimental paradigms. Finally, our model predicted how the available CBP and protein *P*, necessary for promoting potentiation within the local dendritic vicinity, were shared between inputs, leading to the variable changes in spine structure, both at the stimulated and non-stimulated spines.

### Relation to previous studies on clustered stimulation

Previous studies have employed similar experimental techniques to explore heterosynaptic interactions on single stretches of dendrites (Harvey and Svoboda, 2007; Oh et al., 2015; Tong et al., 2021) but showed a variety of seemingly incompatible plasticity outcomes. The differences can be understood using insights from our model. Moreover, many prior sLTP studies using glutamate uncaging have involved potentiating a single isolated spine on a dendrite, thereby likely minimizing the impact on heterosynaptic inputs (for example Matsuzaki et al., 2004; Harvey and Svoboda, 2007). We consistently find no significant heterosynaptic dynamics when we target a single spine to elicit potentiation.

A small group of studies have employed multiple spine uncaging on dendritic segments. For example, in the study by Oh et al. (2015), eliciting LTP in a group of clustered spines resulted in heterosynaptic depression of an unstimulated spine inside the cluster, which was dependent on calcineurin activity. Whereas this finding differs from our current result, where we see potentiation of inside cluster spines, the apParént disagreement can be explained by differences in stimulation paradigms. Oh et al. (2015) stimulated each LTP spine on its own, one by one, in turn, a condition unlikely to generate a substantial dendritic Ca^2+^ transient. Moreover, the time taken to complete stimulating all the targeted spines would have been longer (at least 6 min) than the 60 sec in the current study. Such a major difference in the activity pattern likely engages different endogenous dendritic mechanisms. Govindarajan et al. (2011) stimulated groups of spines either on a single dendrite or across a pair of dendritic branches and measured the dependence of homosynaptic long-lasting LTP (i.e., L-LTP) on a local supply of freshly translated protein. In this study, the strength, frequency, and number of pulses received by each homosynaptic spine were varied between the single spine condition and the multiple spine condition, unlike in the current study where the glutamate uncaging stimulation condition for each target was kept constant irrespective of the number of stimulated spines. Further work by Chirillo et al. (2019) supported the idea that the amount of locally available Ca^2+^ is controlled by clusters of simultaneously active synapses.

Finally, previous work by Tong et al. (2021) has demonstrated a mixture of heterosynaptic potentiation and depression following the simultaneous potentiation of groups of spines on single dendritic branches. Again, the stimulation parameters’ differences may explain differences in the measured outcome. Tong et al. (2021) used a whole-cell patch-clamp to depolarize the postsynaptic neuron to 0mV for one minute in the presence of extracellular *Mg^2+^* to promote NMDAR activation whilst uncaging glutamate onto the homosynaptic spines to elicit structural plasticity. This likely initiated broader and more extensive signalling in the dendrite/neuron, whereas, in the current study, the experiments were performed in nominally magnesium-free aCSF without deliberate depolarization of the postsynaptic cell membrane.

It is unclear which protocol best represents the *in vivo* condition where it is highly likely that multiple inputs may be synchronously active on a single dendrite during behaviour and, in turn, require the postsynaptic machinery to recognize this local spatio-temporal synchrony and respond accordingly. Regardless, our study attempted to identify the minimal components of molecular signalling machinery that can represent the behaviour of active and inactive spines along stretches of dendrites.

### Local versus global effects

On the spatial and temporal scales investigated, we found that the spines sharing a short region of dendrite appeared not to be competing for a limited resource. However, this result does not rule out changes that may be occurring elsewhere in the neuron, outside the imaging ROI, nor can it account for plasticity events that may act to rebalance the network on longer timescales than explored here. Presumably, the clustered potentiation induced in these experiments must have upper limits (as demonstrated in the 15 spine condition), and homeostatic mechanisms that normalize neuronal activity levels (such as synaptic scaling of AMPARs) may operate over slower timescales to be monitored in the current study. Following these dendrites on longer timescales (hours to days) may reveal additional patterns of homosynaptic- and heterosynaptic plasticity that are not apParént here.

The stimulation paradigm we considered did not involve a combination of pre- and post-stimulation and was, therefore, non-Hebbian; we stimulated only presynaptically by uncaging glutamate, thereby inducing sLTP directly. We observed that a single stimulation induced a small amount of sLTP. However, when 3, 7, or 15 spines were stimulated, spine cooperativity emerged, which is consistent with a Hebbian learning component (e.g., via increased voltage at the postsynaptic dendritic segment, which is shared by all synapses). This is in line with previous work, such as Tong et al. (2021), which reported that local glutamate uncaging with cell-wide depolarization and, for groups of 7 spines, homosynaptic inputs lead to robust structural potentiation. In contrast, nearby heterosynaptic inputs shrank in spine size. Building on our findings and combining pre- and postsynaptic stimulation in future experiments could help dissect the interaction of Hebbian and non-Hebbian components across time and dendritic space by studying the sLTP/sLTP by stimulating a controlled number of spines by varying the timing of the glutamate uncaging relative to the timing of a depolarizing current into the dendrite (Yagishita et al., 2014).

Additionally, regarding the sum total growth of estimated spine volumes we have observed, we note that we cannot exclude the local synthesis of proteins. The results obtained within the studied spatial and temporal scales are consistent with other studies. This effect may require additional consideration in revising the model for a future study as the total stores shared among spines may not be constant in time or space, as is assumed here.

Finally, we discuss the global effects that may emerge tens to hundreds of micrometres away from the stimulation site. Our experimental paradigm is restricted to a maximum length of 33 *μ*m. Therefore, we can only comment on the effects observed within this range. In figure S1, we specifically studied the spines outside the stimulated cluster and observed LTD, whereby the LTD amplitude declined with a growing distance from the stimulated cluster. This collective LTD observed in the distant spines was measurable but close to the significance threshold when considering individual spines and their relative change post-stimulation. Thus, it had a smaller amplitude than the plasticity we measured at the stimulated sites and their immediate vicinity.

### Presynaptic identity

Our current study focused entirely on monitoring dendritic spine strength using the intensity of a fluorescent protein (GFP) as a readout. This is a well-used and validated approach, both *in vitro* and *in vivo*, and for example, has been used to demonstrate sLTP associated with behavioural task acquisition (Matsuzaki et al., 2004; Hayashi-Takagi et al., 2015). Single-cell expression of the fluorophore allows excellent signal-to-noise readout of GFP spine signals and the ease of imaging large populations of inputs over time. However, the method does not give information about the presynaptic identity of the axons arriving onto the spines. Our study used apical oblique dendrites of CA1 neurons, and thus the incoming axons were likely part of the Schaffer collaterals originating in CA3 (Sorra and Harris, 1993). These inputs have been shown to make multiple contacts onto single CA1 neurons (Druckmann et al., 2014), including onto single dendritic branches.

It would be of interest to determine the plasticity rules when a cluster of dendritic spines sharing the same presynaptic input is potentiated and how well such rules are matched to spine behaviour following plasticity induction with synchronous glutamate uncaging that bypasses communication with the presynaptic terminals. In our study, glutamate uncaging of smaller cluster sizes (3 and 7 spines) could reflect a single input, or a pair of inputs, being potentiated together. We note that a related study (Bartol Jr et al., 2015) showed that same-dendrite same-axon pairs are rare in the middle of s. radiatum and hence the presynaptic axon could not be conclusively linked to cluster coordination. In contrast, these same-dendrite same-axon pairs were more common in the distal tufts of CA1 and might lead to fruitful future study (Bloss et al., 2018). Further work is required to determine if individual inputs are recognized and privileged from a postsynaptic point of view.

### Computational models of heterosynaptic effects

With the introduction of the protein *P*, our model aimed to retain the essence of the Ca^2+^ amplitude hypothesis, which states that the peak Ca^2+^ determines the direction of plasticity (either sLTP or sLTD) (Lisman, 1989, 2001; Evans and Blackwell, 2015). While our model did not directly include a Ca^2+^ term, the model indirectly accounts for Ca^2+^ changes through the CBP variable, which drives the *P* dynamics. Specifically, instead of Ca^2+^, a low *P* elevation results in sLTD, while a higher elevation resulted in sLTP in our model.

Experimental studies that motivate this hypothesis include the work by Artola et al. (1990) showing that post-synaptic depolarization level could affect the direction of plasticity while the stimulation frequency was kept constant. Cho et al. (2001) showed that directly lowering the extracellular Ca^2+^ concentration turned sLTP into sLTD. In addition, using intracellular uncaging of Ca^2+^, (Yang et al., 1999) reported that a brief but high elevation of Ca^2+^ resulted in sLTP, whereas a longer but slight rise in Ca^2+^ triggered sLTD. To simulate this effect, several models have been proposed that manipulate the Ca^2+^ amplitude and evaluate either frequency-based (Artola et al., 1990; Hansel et al., 1996), STDP-based plasticity rules (for data, see Graupner and Brunel, 2010) or both (Shouval et al., 2002; Bush and Jin, 2012; Helias et al., 2008). Several important predictions have arisen from these models, including the supralinear relationship between stimulation frequency and calcium amplitude (Gamble and Koch, 1987) or Shouval et al. (2002) whose model was able to fit the frequency-based plasticity curve and partly match a spike timing (STDP) based plasticity response.

Interestingly, previous studies have also demonstrated that relying solely on calcium amplitude will always lead to discrepancies with experimental results, e.g. Rubin et al. (2005), showed that any such pure calcium-driven LTP model will always exhibit a second sLTD window at long positive pre-post timing intervals. Nevertheless, several studies addressing this question did not find evidence of this phenomenon and showed no LTD during positive pre-post intervals (Markram et al., 1997; Feldman, 2000; Pawlak and Kerr, 2008). While some experiments did exhibit this second sLTD window, including work by Nishiyama et al. (2000),Wittenberg and Wang (2006), and Inglebert et al. (2020). The seeming conflict indicates that variability in the stimulation protocols, including variable timing and numbers of stimulated sites, can lead to variable plasticity outcomes and combining the action of a slow protein component, as we have introduced in our work, could help understand the mechanisms and identify the stimulation protocols that result in similar sLTP/sLTD outcomes.

Other, more recent models looking beyond the amplitude hypothesis include work by Ebner et al. (2019) that modelled cooperative plasticity exists across the dendritic tree as well as within single branches on a scale of milliseconds, and Triesch et al. (2018) that used a finite pool of receptors to show a variety of heterosynaptic behaviours as a consequence of competitive effects.

In building our model, we implemented a fast Ca^2+^ mediated component and a slower protein component. One of the reasons we did not build directly on the models of Castellani et al. (2001) or Shouval et al. (2002), which also describe plasticity across spines, is the short times of seconds to tens of seconds these models are built to replicate. While in our work, we aimed to capture spine plasticity effects evolving on the time scale of minutes to tens of minutes. Our model is motivated by prior models that aimed to cover different time scales while simplifying the dynamics to two main actors. Examples of such models include the work of Clopath et al. (2008), which used the concept of synaptic tagging to study early and late LTP, as well as Ziegler et al. (2015), which developed a three-layered model of synaptic consolidation through the interaction of the synaptic efficacy with a scaffolding variable by a read-write process mediated by a tagging-related variable and the model by Fusi et al. (2005) which considered discrete states to study the larger idea of how memory retention scales with the number of synapses.

### Biological candidates for *P* and CBP

Up to this point, we have not speculated on the exact nature of the CBPs and *P*. However, several candidates have been previously studied that might represent these quantities. We stress that CBPs and *P* may not necessarily be restricted to single proteins but may represent a group of proteins that act collectively to achieve the results seen in the experiments.

In our experiments, we have presented results pertaining to the inhibition of calcineurin and CaMKII. Calcineurin inhibition has previously been seen to prevent heterosynaptic sLTD without affecting homosynaptic sLTP within a spatial extent of 4*μ*m of the activation site (Oh et al., 2015; Tong et al., 2021), whilst inhibition of CaMKII leads to deficits in sLTP (Silva et al., 1992) and heterosynaptic sLTP (Tong et al., 2021). Other candidate molecules for *P* include Inositol trisphosphate receptors (IP3Rs) which have a similar effect on heterosynaptic sLTP as calcineurin (Oh et al., 2015), h-Ras which has been seen to lower the threshold for neighbouring spine sLTP induction (Harvey et al., 2008) or RhoA which, following synaptic activity, exits the spine and diffuses along the dendrite (Murakoshi et al., 2011).

Candidates for CBP include, Ca^2+^ which is a key factor for synaptic plasticity (Evans and Blackwell, 2015), the ARC gene product, which acts in the form of slow homosynaptic sLTP followed by heterosynaptic sLTD (El-Boustani et al., 2018) or NO synthase that produces nitric oxide, which in turn facilitates sLTP and affects synaptic strengths of neighbouring neurons (Schuman and Madison, 1994; Padamsey et al., 2017).

Designing experiments with these quantities in mind could reveal more about the nature and composition of both the CBP and *P*.

### Model insights into multi-spine sLTP

We have introduced a model that is based on fast Ca^2+^ action fueling the activation of a long-lasting protein *P* whose concentration ultimately drives the sLTP vs sLTD plasticity decision at the individual spines. Building on the previously introduced Ca^2+^ levels hypothesis (Lisman, 1989; Castellani et al., 2001), we could explain experimental outcomes across different spatio-temporal combinations of induction sites while retaining the basic features of homo- and heterosynaptic plasticity. Our model, which is continuous in time, allowed for the study of both short-term and long-term saturation effects of plasticity, as observed by the parameter fits for the experimental results across different experimental conditions. Interestingly, our model predicted the experimentally observed spine change across different experimental conditions using only two dynamic variables, CBP and *P*. Additionally, we were able to identify a set of invariant parameters that can predict the plasticity response across any number of stimulation events, and we cross-validated these parameters using our experiment with 15 stimulation sites.

We also found that the inside neighbour spine response could be explained only by varying the initially available CBP and *P*. Interestingly, even though our model had only two dynamic variables and averaged over the spatial dynamics and operated with an effective description of the LTP/LTD boundary, it could accurately describe the plasticity and its time course across different numbers of stimulated spines. These insights will allow the model to be employed in spiking network simulations and give insight into the circuit-level consequences of the experimentally observed multi-spine plasticity rules.

## Methods

### Experimental

#### Preparation of organotypic hippocampal slice culture

Organotypic hippocampal slices were preParéd as previously reported (Stoppini et al., 1991). Briefly, hippocampi of postnatal day P6-7 Wistar rat pups (Nihon SLC) were isolated and cut into 350 *μ*m-thick transverse slices on a McIlwain tissue chopper (Mickle Laboratory Engineering Co. sLTD. and Cavey Laboratory Engineering Co. sLTD.). Slices were transferred onto cell culture inserts (0.4 mm pore size, Merck Millipore) and placed in a 6-well cell culture plate filled with 1 ml/well of culture media containing 50% Minimum Essential Medium (MEM, Thermo Fisher Scientific), 23% Earle’s Balanced Salt Solution, 25% horse serum (Thermo Fisher Scientific), and 36 mM D-glucose. Slices were maintained at 35°C in 5% CO_2_ and used for experiments at 14-18 days in vitro.

During experiments, slices were constantly perfused (1-2 ml/min) with artificial cerebrospinal fluid (aCSF) containing (in mM) 125 NaCl, 2.5 KCl, 26 NaHCO_3_, 1.25 NaH_2_PO_4_, 20 glucose, 2 CaCl_2_ and 4 MNI-glutamate (Tocris). aCSF was continually bubbled with 95%O_2_ and 5%CO_2_, and experiments were carried out at room temperature. For a subset of experiments, calcineurin or CaMKII was inhibited with FK506 (2*μ*M, Tocris) or myristoylated Autocamtide-2-related inhibitory peptide (AIP, 5*μ*M, Calbiochem) respectively. All animal experiments were approved by the RIKEN Animal Experiments Committee and performed in accordance with the RIKEN rules and guidelines [Animal Experiment Plan Approval no. W2021-2-015(3)].

#### Transfection and imaging of CA1 pyramidal neurons

Organotypic slices were biolistically transfected using a Helios gene gun (Bio-Rad) and used for experiments 48-96 hours later. For structural plasticity experiments, gold particles were coated in a plasmid encoding EGFP. 50*μ*g of EGFP plasmid were coated onto 20-30mg of 1.6*μ*m gold particles. Neurons were imaged at 910nm on a Zeiss LSM 780 confocal laser scanning microscope, and all data were analysed offline.

#### Dendritic spine imaging and glutamate photolysis

Regions of dendrites were chosen by eye for imaging and stimulation. Regions were imaged for a brief baseline period by collecting *z* stacks of the dendritic arbour (512×512, 4× digital zoom for a final frame size of 33.7*μ*m). The z step was 0.5*μ*m. Glutamate was uncaged onto spines lying in the focal z plane using custom-written software at a distance of 0.5*μ*m from the spine head. Medium-sized spines with a clear spine head within the field of view were preferentially targeted for stimulation. MNI-glutamate was photolyzed with a 2-photon laser source (720 nm), and each dendritic spine received a train of 60 pulses of laser light, each 4 msec long, repeated at 1Hz.

For sham experiments, MNI-glutamate was omitted from the aCSF. For groups of stimulated spines, laser pulses were delivered in a quasi-simultaneous fashion in sequence, in which the first spine received a pulse of glutamate (4msec), which was followed by a short delay (<3msec) as the system positioned the laser to the next spine in the sequence. This was repeated for all targeted spines in the stimulated cluster (3, 7, or 15) and the sequence was repeated at 1Hz for 60 cycles.

### Numerical

#### Image Analysis

Estimated spine volumes were obtained from background-subtracted maximum-projected fluorescence images using the integrated pixel intensity (see Chen et al., 2015; Bartol Jr et al., 2015) of an octagonally shaped ROI surrounding the spine head. These values were normalized against the three observed data points immediately preceding the glutamate uncaging. The spine ROI was generated by using a semi-automatic in-house python package that took advantage of the structures of the spines (see the supplemental information for a full list and description) to generate a reproducible ROI. The manual interaction involves a simple clicking on the interior of the spines while the ROI and subsequent measurement are performed automatically. Temporal shifting was corrected using a phase cross-correlation algorithm implemented in SciPy (Guizar-Sicairos et al., 2008). The entirety of this algorithm is designed and written in a user-friendly python package available for download.

Synapses that were partially obscured by the dendrite or overlapped with other spines were omitted from the analysis. A significance test determined the success of an experiment in the form of a z-test, with less than 15% of experiments considered failures. All images shown and used for analysis are maximum-intensity projects of the 3D stacks.

#### Statistical definitions

The spines’ fluorescences obtained from our ROI detection algorithm were normalised against the mean of their pre-stimulation values as follows

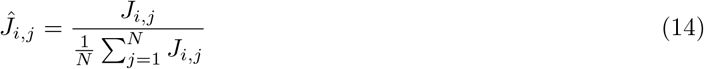

where *J_i,j_* is the fluorescence of spine *i* at snapshot *j* and *N* is the number of pre-stimulation snapshots (in our experiment *N* = 3). Once the spines’ fluorescences were normalised against their baseline, we pooled them across repetitions of the same experiment and cells. All statistics were calculated using this measure. Error bars represent the standard error of the mean, and significance was set at *P* = 0.05 (studentised bootstrap). To comParé different paradigms, Welch’s unequal variances t-test was performed. One asterisk indicates *P* < 0.05.

#### Model fitting algorithm

The values of the parameters *P*_0_, *C*_0_, *α*, *β*_1,2_, *ζ*_1,2_ and *ν_P,D_* of the model (see the equation in figure Fig. 1f) were obtained using a non-linear least-squares approach. The fitting routine included the introduction of a cost function that, when minimised, enforces agreement between the simulated model and observed data points and an iterative update scheme that finds parameters that best represent the experimental data. Mathematically, the cost function can be defined as

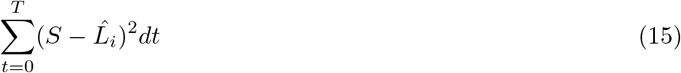

where *T* is the length of the experiment (in our case, 40 minutes) and *L_i_* refers to the averaged response of the spine of type *i* (which can be the stimulated or inside neighbour spines). We note, however, that the experimental snapshots are discrete in time, and the above formulation is continuous. Thus, the cost function needs to be rewritten to reflect the discontinuity as follows

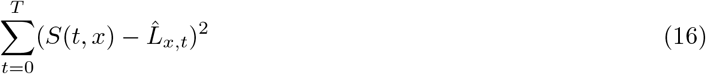

where 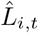 is the spine response at snapshot *t*. The iterative scheme we chose to employ is a gradient-based adjoint approach (Jameson, 1988, or Skene et al., 2021). The adjoint method was chosen, in part, due to its ability to easily and efficiently handle multiple simultaneous optimisation parameters. A numerical solution, implemented in a python code and available on request, supported the optimisation routine to provide feedback on the model dynamics given a set of parameters. The initial parameter fits were obtained with the values obtained from the three-spine paradigm. The quantities that we considered independent across experimental set-ups (*α*, *β*_1,2_, *ζ*_1,2_) were frozen. The other control paradigms were obtained by altering the initial amount of *C* and *P* as well as the *P* and the LTD/LTP peak locations (*ν_P,D_*).

Similarly, these values were fitted in the multi-spine experiments to understand the dynamics of the heterosynaptic spines. Finally, for the experiments where CaMKII and calcineurin blockers were present, only the initial *P*_0_ and the LTP/LTD peak locations (*ν_P/D_*) were changed. We note that the sham experiments were fitted by setting the initial *C*_0_ and *P*_0_ (values inherent to the stimulation process) to 0. For each optimisation, the convergence of obtained parameter fits was ensured by initiating the routine ten separate times with randomly chosen initial parameter configurations.

We considered and fitted only two spine types, the stimulated synapses and spines inside a stimulation site cluster. While we initially considered a spine-specific model, we followed the convention of the field and built an effective model that averages across individual spines and their variable signal-to-noise ratios.

## Data and code availability

Experimental data sets included in the manuscript and the computer code underlying this study can be found in this public github repository https://github.com/meggl23/MultiSpinePlasticity.

## Acknowledgements

This research was supported by University of Bonn Medical Center, University of Mainz Medical Center, ReALity program at the Mainz Medical Center, RIKEN Center for Brain Science, JSPS Core-to-Core Program (JPJSCCA20220007 to YG), and by an add-on fellowship of the Joachim Herz Stiftung (ME). TT and YG thank all our group members for fruitful discussions, and Janko Petkovic for feedback on an earlier version of the manuscript (TT). This project has received funding from the European Research Council (ERC) under the European Union’s Horizon 2020 research and innovation programme (‘MolDynForSyn’, grant agreement No. 945700)

## Supplemental Information

**Figure S1:**
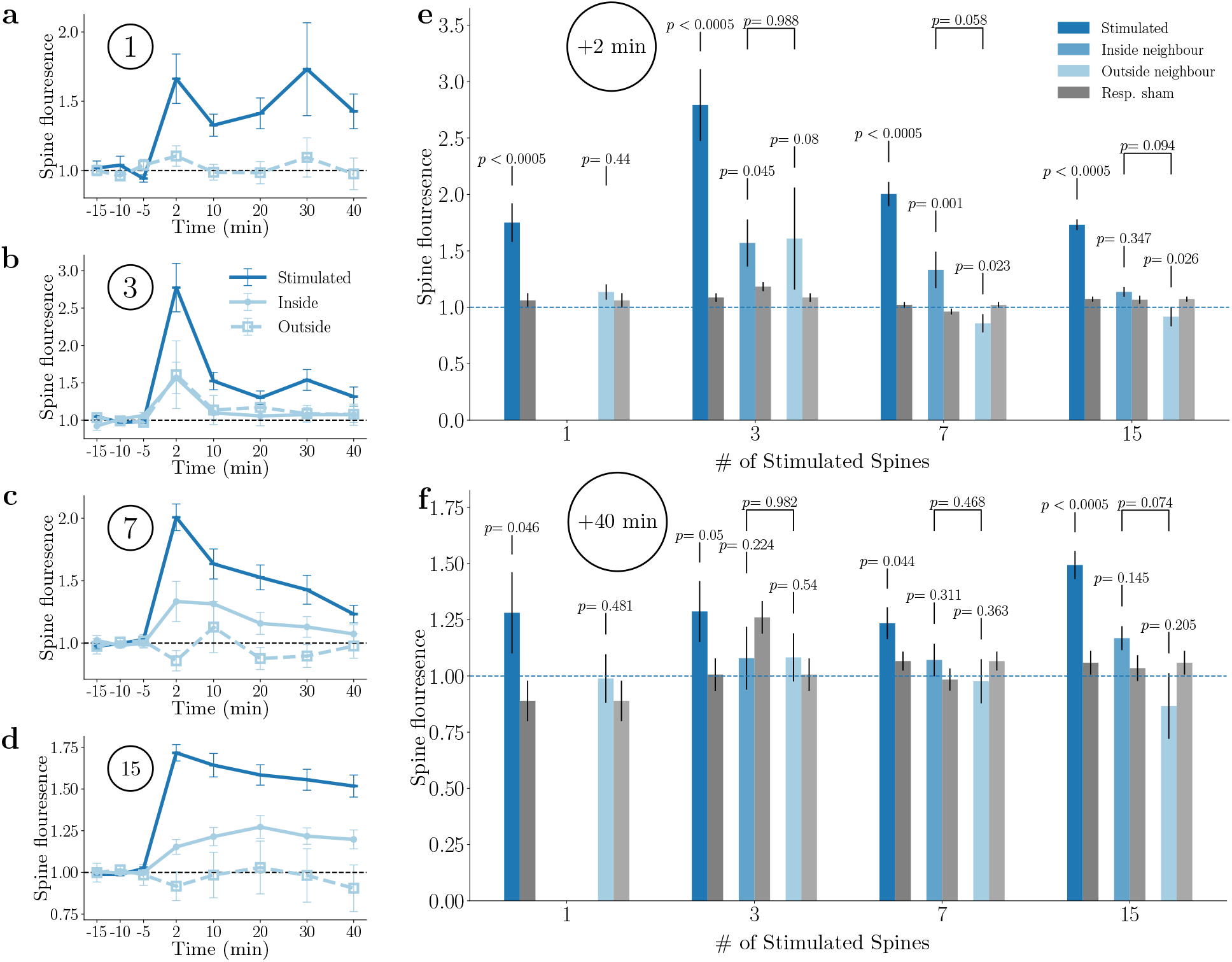
After sLTP induction, stimulated spines and inside neighbour spines both grow while the outside neighbour spines (< 2*μ*m) shrink. **a)**-**d)** Comparative plot of the stimulated, inside neighbour and outside neighbour spines for the single spine, three spine, seven spine and 15 spine paradigm, respectively. **e)** Normalised growth of control conditions of the stimulated, inside neighbour and outside neighbour spines at *t* = 2 minutes. The grey refers to the respective outside neighbours of the sham set. * signifies *p* < 0.05. **f)** Normalised growth of control conditions of the stimulated, inside neighbour and outside neighbour spines at *t* = 40 minutes. The grey refers to the respective outside neighbours of the sham data set.

**Figure S2:**
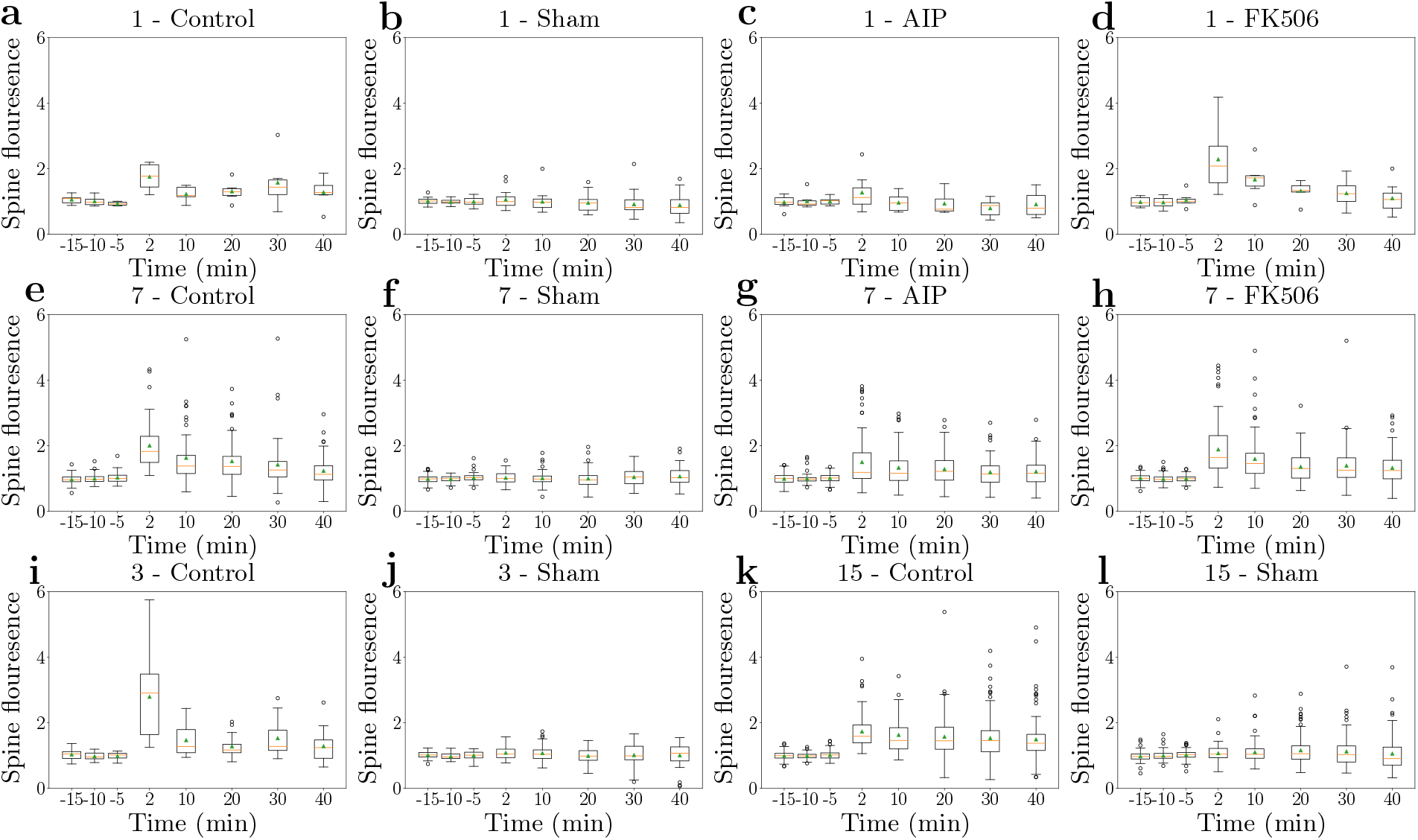
Experimentally recorded normalised statistics of the stimulated spines in experiments with 1, 3, 7 or 15 stimulated spines. **a) - d)** Dynamics of the single spine experiment for the control and sham. Lower to upper quartile values of the data are shown, with a line denoting the median. The whiskers refer to 1.5 times the IQR. Data points outside the whiskers are considered outliers. The green triangle refers to the mean of the data. **e) - f)** Dynamics of the seven spine experiment with the four different paradigms. **i) - j)** Dynamics of the three spine experiment for the control and sham. **k) - l)** Dynamics of the 15 spine experiment for the control and sham.

**Figure S3:**
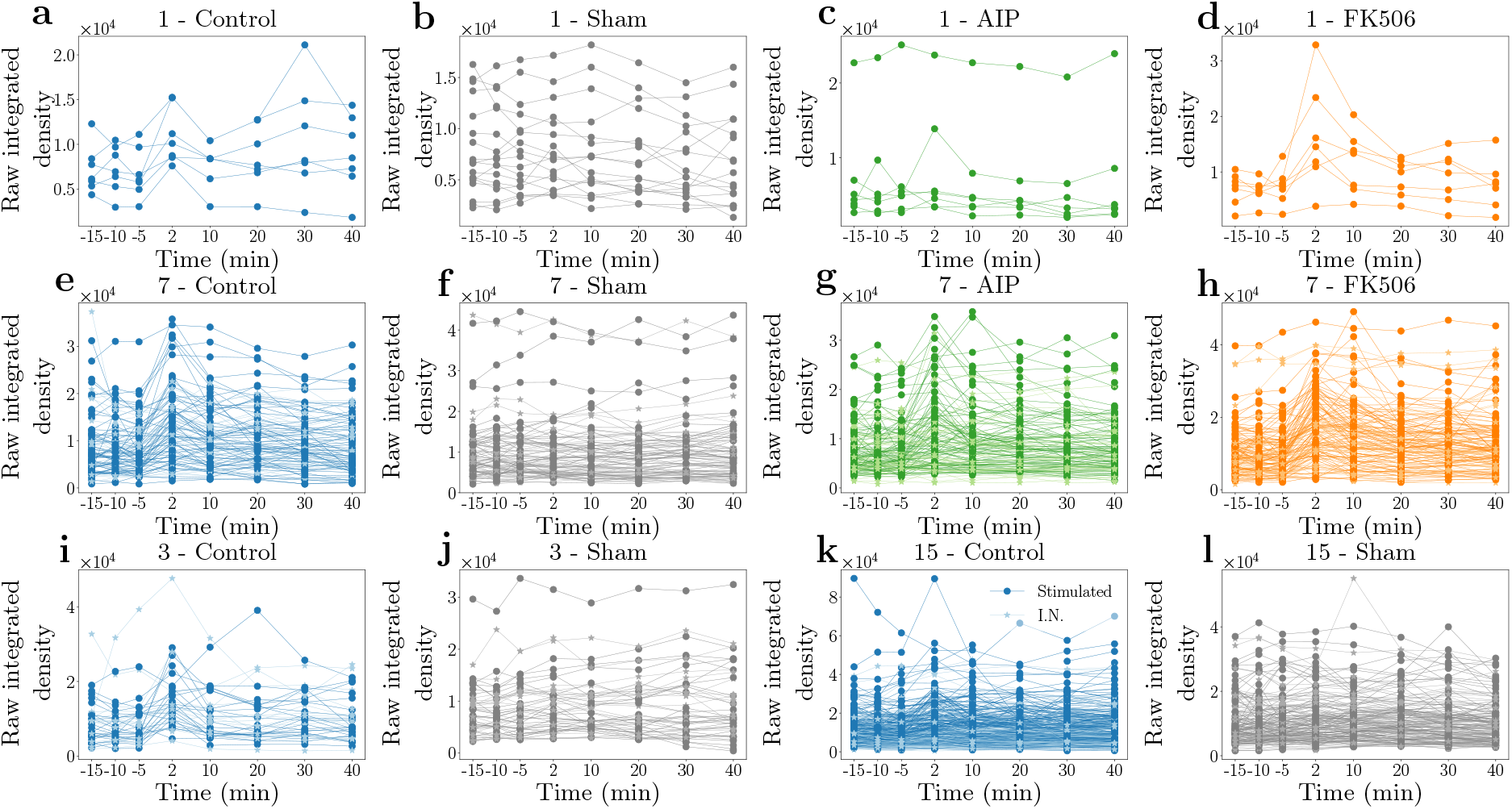
Experimentally recorded unnormalised statistics of the stimulated spines and inside neighbour spines in experiments with 1, 3, 7 or 15 stimulated sites. **a) - d)** Dynamics of the single spine experiment with the four different paradigms. **e) - f)** Dynamics of the seven spine experiment with the four different paradigms. Darker colors refer to stimulated spines and lighter colors to inside neighbour spines. **i) - j)** Dynamics of the three spine experiment for the control and sham. **k) - l)** Dynamics of the 15 spine experiment for the control and sham.

### ROI detection algorithm

To identify a spine ROI, we needed to (*i*) identify the dendrite position and (*ii*) manually select the spine centre. To identify the dendrite position, we clicked on the beginning and end of the dendrite stretch of interest, and a breadth-first search (Moore, 1959) identified the dendritic path. We first needed to determine a putative spine centre to calculate the spine ROI by manually selecting the centre coordinate *x*_0_. Once the spine centre was obtained, the ROI was calculated by stepping outward, pixel by pixel, on eight rays that formed an irregular octagon. Next, our algorithm checked if the following rules were met at each step, and if any of them were broken, a counter (also called a strike) was increased proportionally at each point where a strike occurred. A strike was added if the following conditions were met:

1. **Boundary Rule** If the ROI ray exceeds the boundary of the image, this ends the progression in that direction.
2. **Contour Rule** The image is treated with a Canny edge detection algorithm which calculates edges in an image. Depending on the distance from *x*_0_, encountering such an edge counts as a certain number of strikes.
3. **normalised fluorescence Fall-off Rule** Suppose the normalised fluorescence of the test point becomes one-third of the initial normalised fluorescence of *x*_0_, or the normalised fluorescence falls below four times the background. In that case, the algorithm counts this rule as broken.
4. **Dendrite Rule** We assume that the spine is on average symmetrical and that the user will select the centre of the spine. Therefore, assuming that the initial point *x*_0_ is outside the dendrite, this rule is broken once the test point is closer to the centre of the dendrite than to *x*_0_.
5. **normalised fluorescence Increase Rule** Suppose the normalised fluorescence increases on consecutive steps away from *x*_0_. In that case, we have most probably entered the dendrite, and so after a certain amount of steps, we also consider this a rule break.

Once all rays have stopped, the ROI is drawn by connecting the eight points into an irregular octagon encompassing the spine. We used this octagon ROI to determine the normalised fluorescence quantities and calculate its area. To determine the interior of the ROI, we use a simple test involving the winding number algorithm (Hormann and Agathos, 2001), which sums up the angles subtended by each side of the polygon. If this number is non-zero, the test point is inside the polygon.

### Details on the parameter fitting algorithm

Here, we present the equations that drive the gradient-based adjoint approach (Magri and Juniper, 2014) described in the model analysis section but will omit the derivation thereof as it lies outside the scope of this article. Direct solutions of the gradients are evaluated numerically to acquire the required parameter updates. The first three equations define the adjoint state variables (denoted by the (·)^†^) that deliver the sensitivity information of the model to changes in the parameters. The parameters without the dagger remain the same as those observed in the governing model equations. Finally, 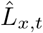 was introduced in the numerical methods section and denotes the averaged spine

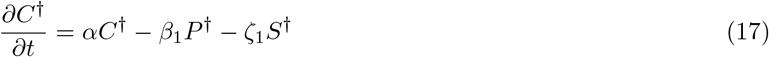

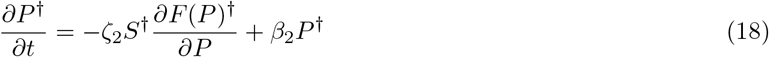

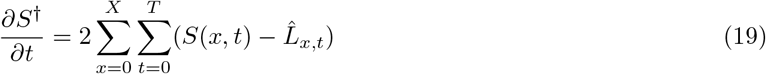

The following equations denote the required calculations involving both the model and adjoint variables to calculate the gradient that will lead to better fits. Each of these gradients is denoted by the variable we wish to change and the superscript †.

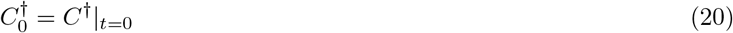

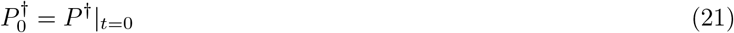

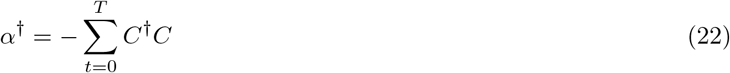

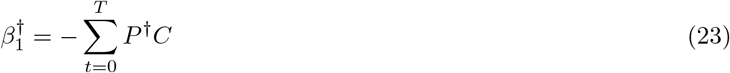

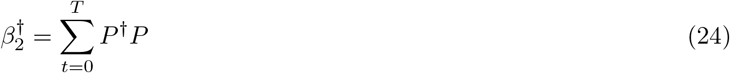

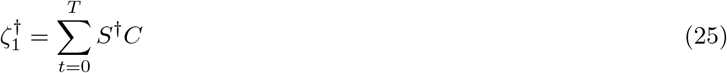

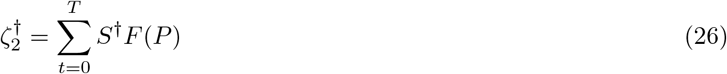

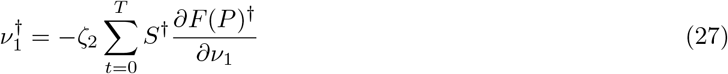

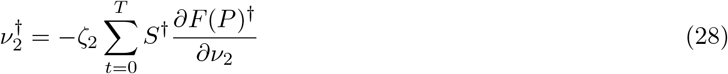

Given these equations, a simple example update step would then be

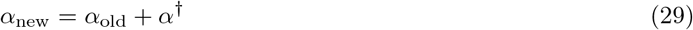

We thus iterate over our parameters until no more change is observed in the update values.

### Parameter optimisation strategy

Here we report the number of experiments and selected spines for each experimental condition. All of these points were acquired through the semi-automatic python tool with manual re-correction in ≈ 5%(81) of the spines.

### Spatial reduction justification

Considering diffusive dynamics of the form with where *α*_1_, *α*_2_ > 0

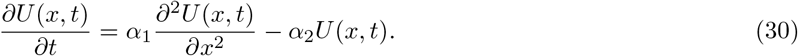

The solution can be separated into a product of a temporal component *T*(*t*) and spatial component *X*(*x*)

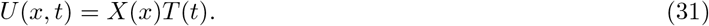

Substituting this into the equation above and rearranging the terms to

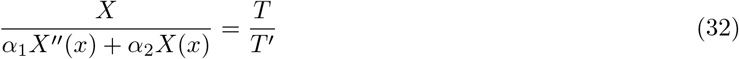

Given that the left-hand side is only dependent on *x* and the right-hand side only depends on *t*, we can see that these terms need to be constant, which we define as *λ*. This leads us to a pair of tractable ordinary differential equations

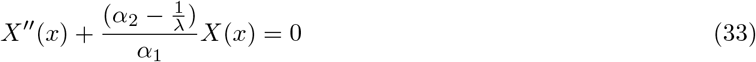

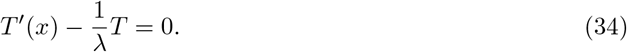

Defining 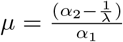, we obtain

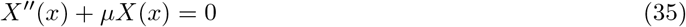

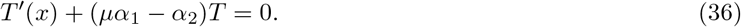

The solution for the two quantities is:

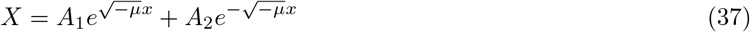

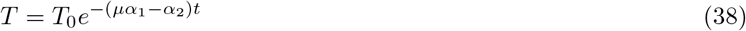

where *A*_1_ and *A*_2_ arise from boundary conditions at the spatial boundary of the considered domain and at the starting time *t* = *T*_0_. Thus considering the single spine model, i.e., *x* = 0, leads to

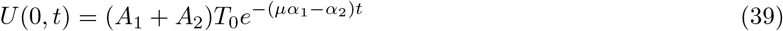

Therefore, we can introduce the terms 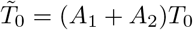 and 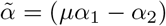 to note that up to multiplicative constants, we the above system decays exponentially like the single spine mode at *x* = 0, i.e.,

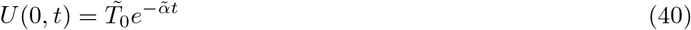

## Notes

### Competing Interest Statement

The authors have declared no competing interest.

### Summary of Updates

Introduction and results section updated to better introduce model design.

## References

Alain Artola, S Bröcher, and Wolf Singer. Different voltage-dependent thresholds for inducing long-term depression and long-term potentiation in slices of rat visual cortex. Nature, 347(6288):69–72, 1990.

Thomas M Bartol Jr, Cailey Bromer, Justin Kinney, Michael A Chirillo, Jennifer N Bourne, Kristen M Harris, and Terrence J Sejnowski. Nanoconnectomic upper bound on the variability of synaptic plasticity. Elife, 4: e10778, 2015.

Elie L Bienenstock, Leon N Cooper, and Paul W Munro. Theory for the development of neuron selectivity: orientation specificity and binocular interaction in visual cortex. Journal of Neuroscience, 2(1):32–48, 1982.

Erik B Bloss, Mark S Cembrowski, Bill Karsh, Jennifer Colonell, Richard D Fetter, and Nelson Spruston. Single excitatory axons form clustered synapses onto ca1 pyramidal cell dendrites. Nature neuroscience, 21 (3):353–363, 2018.

Tobias Bonhoeffer, Volker Staiger, and AMHJ Aertsen. Synaptic plasticity in rat hippocampal slice cultures: local” hebbian” conjunction of pre-and postsynaptic stimulation leads to distributed synaptic enhancement. Proceedings of the National Academy of Sciences, 86(20):8113–8117, 1989.

Miquel Bosch, Jorge Castro, Takeo Saneyoshi, Hitomi Matsuno, Mriganka Sur, and Yasunori Hayashi. Structural and molecular remodeling of dendritic spine substructures during long-term potentiation. Neuron, 82(2):444–459, 2014.

Tiago Branco and Michael Häusser. Synaptic integration gradients in single cortical pyramidal cell dendrites. Neuron, 69(5):885–892, 2011.

Daniel Bush and Yaochu Jin. Calcium control of triphasic hippocampal stdp. Journal of computational neuroscience, 33(3):495–514, 2012.

Pico Caroni, Flavio Donato, and Dominique Muller. Structural plasticity upon learning: regulation and functions. Nature Reviews Neuroscience, 13(7):478–490, 2012.

Gastone C Castellani, Elizabeth M Quinlan, Leon N Cooper, and Harel Z Shouval. A biophysical model of bidirectional synaptic plasticity: dependence on ampa and nmda receptors. Proceedings of the National Academy of Sciences, 98(22):12772–12777, 2001.

Thomas E Chater and Yukiko Goda. My neighbour hetero—deconstructing the mechanisms underlying heterosynaptic plasticity. Current Opinion in Neurobiology, 67:106–114, 2021.

Xiaobing Chen, Jonathan M Levy, Austin Hou, Christine Winters, Rita Azzam, Alioscka A Sousa, Richard D Leapman, Roger A Nicoll, and Thomas S Reese. Psd-95 family maguks are essential for anchoring ampa and nmda receptor complexes at the postsynaptic density. Proceedings of the National Academy of Sciences, 112 (50):E6983–E6992, 2015.

Marina E Chicurel and Kristen M Harris. Three-dimensional analysis of the structure and composition of ca3 branched dendritic spines and their synaptic relationships with mossy fiber boutons in the rat hippocampus. Journal of comparative neurology, 325(2):169–182, 1992.

Michael A Chirillo, Mikayla S Waters, Laurence F Lindsey, Jennifer N Bourne, and Kristen M Harris. Local resources of polyribosomes and ser promote synapse enlargement and spine clustering after long-term potentiation in adult rat hippocampus. Scientific reports, 9(1):1–14, 2019.

K Cho, John Patrick Aggleton, MW Brown, and ZI Bashir. An experimental test of the role of postsynaptic calcium levels in determining synaptic strength using perirhinal cortex of rat. The Journal of physiology, 532 (2):459–466, 2001.

Claudia Clopath, Lorric Ziegler, Eleni Vasilaki, Lars Buüsing, and Wulfram Gerstner. Tag-trigger-consolidation: a model of early and late long-term-potentiation and depression. PLoS computational biology, 4(12):e1000248, 2008.

Shaul Druckmann, Linqing Feng, Bokyoung Lee, Chaehyun Yook, Ting Zhao, Jeffrey C Magee, and Jinhyun Kim. Structured synaptic connectivity between hippocampal regions. Neuron, 81(3):629–640, 2014.

Francis J Dumont. Fk506, an immunosuppressant targeting calcineurin function. Current medicinal chemistry, 7(7):731–748, 2000.

Christian Ebner, Claudia Clopath, Peter Jedlicka, and Hermann Cuntz. Unifying long-term plasticity rules for excitatory synapses by modeling dendrites of cortical pyramidal neurons. Cell reports, 29(13):4295–4307, 2019.

Sami El-Boustani, Jacque PK Ip, Vincent Breton-Provencher, Graham W Knott, Hiroyuki Okuno, Haruhiko Bito, and Mriganka Sur. Locally coordinated synaptic plasticity of visual cortex neurons in vivo. Science, 360(6395):1349–1354, 2018.

Florian Engert and Tobias Bonhoeffer. Synapse specificity of long-term potentiation breaks down at short distances. Nature, 388(6639):279–284, 1997.

RC Evan and KT Blackwell. Calcium: amplitude, duration, or location? The Biological Bulletin, 228(1):75–83, 2015.

Daniel E Feldman. Timing-based ltp and ltd at vertical inputs to layer ii/iii pyramidal cells in rat barrel cortex. Neuron, 27(1):45–56, 2000.

Reiko Maki Fitzsimonds, Hong-jun Song, and Mu-ming Poo. Propagation of activity-dependent synaptic depression in simple neural networks. Nature, 388(6641):439–448, 1997.

Yombe Fonkeu, Nataliya Kraynyukova, Anne-Sophie Hafner, Lisa Kochen, Fabio Sartori, Erin M. Schuman, and Tatjana Tchumatchenko. How mRNA localization and protein synthesis sites influence dendritic protein distribution and dynamics. Neuron, 103(6):1109–1122.e7, sep 2019. doi: 10.1016/j.neuron.2019.06.022.

Min Fu, Xinzhu Yu, Ju Lu, and Yi Zuo. Repetitive motor learning induces coordinated formation of clustered dendritic spines in vivo. Nature, 483(7387):92–95, 2012.

Hajime Fujii, Masatoshi Inoue, Hiroyuki Okuno, Yoshikazu Sano, Sayaka Takemoto-Kimura, Kazuo Kitamura, Masanobu Kano, and Haruhiko Bito. Nonlinear decoding and asymmetric representation of neuronal input information by camkiiα and calcineurin. Cell reports, 3(4):978–987, 2013.

Stefano Fusi, Patrick J Drew, and Larry F Abbott. Cascade models of synaptically stored memories. Neuron, 45(4):599–611, 2005.

Edward Gamble and Christof Koch. The dynamics of free calcium in dendritic spines in response to repetitive synaptic input. Science, 236(4806):1311–1315, 1987.

S Glazewski, KP Giese, A Silva, and K Fox. The role of α-camkii autophosphorylation in neocortical experiencedependent plasticity. Nature neuroscience, 3(9):911–918, 2000.

Arvind Govindarajan, Raymond J Kelleher, and Susumu Tonegawa. A clustered plasticity model of long-term memory engrams. Nature Reviews Neuroscience, 7(7):575–583, 2006.

Arvind Govindarajan, Inbal Israely, Shu-Ying Huang, and Susumu Tonegawa. The dendritic branch is the preferred integrative unit for protein synthesis-dependent ltp. Neuron, 69(1):132–146, 2011.

Michael Graupner and Nicolas Brunel. Mechanisms of induction and maintenance of spike-timing dependent plasticity in biophysical synapse models. Frontiers in computational neuroscience, 4:136, 2010.

Manuel Guizar-Sicairos, Samuel T Thurman, and James R Fienup. Efficient subpixel image registration algorithms. Optics letters, 33(2):156–158, 2008.

C Hansel, A Artola, and W Singer. Different threshold levels of postsynaptic [ca2+] i have to be reached to induce ltp and ltd in neocortical pyramidal cells. Journal of Physiology-Paris, 90(5-6):317–319, 1996.

Christopher D Harvey and Karel Svoboda. Locally dynamic synaptic learning rules in pyramidal neuron dendrites. Nature, 450(7173):1195–1200, 2007.

Christopher D Harvey, Ryohei Yasuda, Haining Zhong, and Karel Svoboda. The spread of ras activity triggered by activation of a single dendritic spine. Science, 321(5885):136–140, 2008.

Akiko Hayashi-Takagi, Sho Yagishita, Mayumi Nakamura, Fukutoshi Shirai, Yi I Wu, Amanda L Loshbaugh, Brian Kuhlman, Klaus M Hahn, and Haruo Kasai. Labelling and optical erasure of synaptic memory traces in the motor cortex. Nature, 525(7569):333–338, 2015.

Moritz Helias, Stefan Rotter, Marc-Oliver Gewaltig, and Markus Diesmann. Structural plasticity controlled by calcium based correlation detection. Frontiers in Computational Neuroscience, 2:7, 2008.

Takao K Hensch. Critical period plasticity in local cortical circuits. Nature Reviews Neuroscience, 6(11):877–888, 2005.

Kai Hormann and Alexander Agathos. The point in polygon problem for arbitrary polygons. Computational geometry, 20(3):131–144, 2001.

Yanis Inglebert, Johnatan Aljadeff, Nicolas Brunel, and Dominique Debanne. Synaptic plasticity rules with physiological calcium levels. Proceedings of the National Academy of Sciences, 117(52):33639–33648, 2020.

Atsuhiko Ishida, Isamu Kameshita, Sachiko Okuno, Takako Kitani, and Hitoshi Fujisawa. A novel highly specific and potent inhibitor of calmodulin-dependent protein kinase ii. Biochemical and biophysical research communications, 212(3):806–812, 1995.

David B Jaffe, Daniel Johnston, Nechama Lasser-Ross, John E Lisman, Hiroyoshi Miyakawa, and William N Ross. The spread of na+ spikes determines the pattern of dendritic ca2+ entry into hippocampal neurons. Nature, 357(6375):244–246, 1992.

A. Jameson. Aerodynamic design via control theory. J. Sci. Comp., 3(3):233–260, 1988.

Narayanan Kasthuri, Kenneth Jeffrey Hayworth, Daniel Raimund Berger, Richard Lee Schalek, José Angel Conchello, Seymour Knowles-Barley, Dongil Lee, Amelio Vázquez-Reina, Verena Kaynig, Thouis Raymond Jones, et al. Saturated reconstruction of a volume of neocortex. Cell, 162(3):648–661, 2015.

Kevin FH Lee, Cary Soares, Jean-Philippe Thivierge, and Jean-Claude Béïque. Correlated synaptic inputs drive dendritic calcium amplification and cooperative plasticity during clustered synapse development. Neuron, 89 (4):784–799, 2016.

Seok-Jin R Lee, Yasmin Escobedo-Lozoya, Erzsebet M Szatmari, and Ryohei Yasuda. Activation of camkii in single dendritic spines during long-term potentiation. Nature, 458(7236):299–304, 2009.

Mathieu Letellier, Florian Levet, Olivier Thoumine, and Yukiko Goda. Differential role of pre-and postsynaptic neurons in the activity-dependent control of synaptic strengths across dendrites. PLoS biology, 17(6):e2006223, 2019.

John Lisman. A mechanism for the hebb and the anti-hebb processes underlying learning and memory. Proceedings of the National Academy of Sciences, 86(23):9574–9578, 1989.

John E Lisman. Three ca2+ levels affect plasticity differently: the ltp zone, the ltd zone and no man’s land. The Journal of physiology, 532(Pt 2):285, 2001.

Gary S Lynch, Thomas Dunwiddie, and Valentin Gribkoff. Heterosynaptic depression: a postsynaptic correlate of long-term potentiation. Nature, 266(5604):737–739, 1977.

Luca Magri and Matthew P Juniper. Adjoint-based linear analysis in reduced-order thermo-acoustic models. International Journal of Spray and Combustion Dynamics, 6(3):225–246, 2014.

Guy Major, Alon Polsky, Winfried Denk, Jackie Schiller, and David W Tank. Spatiotemporally graded nmda spike/plateau potentials in basal dendrites of neocortical pyramidal neurons. Journal of neurophysiology, 99 (5):2584–2601, 2008.

Hiroshi Makino and Roberto Malinow. Compartmentalized versus global synaptic plasticity on dendrites controlled by experience. Neuron, 72(6):1001–1011, 2011. doi: https://doi.org/10.1016/j.neuron.2011.09.036.

Henry Markram, Joachim Lubke, Michael Frotscher, and Bert Sakmann. Regulation of synaptic efficacy by coincidence of postsynaptic aps and epsps. Science, 275(5297):213–215, 1997.

Masanori Matsuzaki, Naoki Honkura, Graham CR Ellis-Davies, and Haruo Kasai. Structural basis of long-term potentiation in single dendritic spines. Nature, 429(6993):761–766, 2004.

Stephen G Miller and Mary B Kennedy. Regulation of brain type ii ca2+ calmodulin-dependent protein kinase by autophosphorylation: A ca2+-triggered molecular switch. Cell, 44(6):861–870, 1986.

Edward F Moore. The shortest path through a maze. In Proc. Int. Symp. Switching Theory, 1959, pages 285–292, 1959.

Rosel M Mulkey and Robert C Malenka. Mechanisms underlying induction of homosynaptic long-term depression in area ca1 of the hippocampus. Neuron, 9(5):967–975, 1992.

Hideji Murakoshi, Hong Wang, and Ryohei Yasuda. Local, persistent activation of rho gtpases during plasticity of single dendritic spines. Nature, 472(7341):100–104, 2011.

Thomas Nevian and Bert Sakmann. Spine ca2+ signaling in spike-timing-dependent plasticity. Journal of Neuroscience, 26(43):11001–11013, 2006.

Makoto Nishiyama, Kyonsoo Hong, Katsuhiko Mikoshiba, Mu-Ming Poo, and Kunio Kato. Calcium stores regulate the polarity and input specificity of synaptic modification. Nature, 408(6812):584–588, 2000.

Won Chan Oh, Laxmi Kumar Parajuli, and Karen Zito. Heterosynaptic structural plasticity on local dendritic segments of hippocampal ca1 neurons. Cell reports, 10(2):162–169, 2015.

Yannan Ouyang, David Kantor, Kristen M Harris, Erin M Schuman, and Mary B Kennedy. Visualization of the distribution of autophosphorylated calcium/calmodulin-dependent protein kinase ii after tetanic stimulation in the ca1 area of the hippocampus. Journal of Neuroscience, 17(14):5416–5427, 1997.

Zahid Padamsey, Lindsay McGuinness, Scott J Bardo, Marcia Reinhart, Rudi Tong, Anne Hedegaard, Michael L Hart, and Nigel J Emptage. Activity-dependent exocytosis of lysosomes regulates the structural plasticity of dendritic spines. Neuron, 93(1):132–146, 2017.

Verena Pawlak and Jason ND Kerr. Dopamine receptor activation is required for corticostriatal spike-timing-dependent plasticity. Journal of Neuroscience, 28(10):2435–2446, 2008.

Roger L Redondo and Richard GM Morris. Making memories last: the synaptic tagging and capture hypothesis. Nature Reviews Neuroscience, 12(1):17–30, 2011.

Jacqueline Rose, Shan-Xue Jin, and Ann Marie Craig. Heterosynaptic molecular dynamics: locally induced propagating synaptic accumulation of cam kinase ii. Neuron, 61(3):351–358, 2009.

Sébastien Royer and Denis Paré. Conservation of total synaptic weight through balanced synaptic depression and potentiation. Nature, 422(6931):518–522, 2003.

Jonathan E Rubin, Richard C Gerkin, Guo-Qiang Bi, and Carson C Chow. Calcium time course as a signal for spike-timing–dependent plasticity. Journal of neurophysiology, 93(5):2600–2613, 2005.

Jackie Schiller, Guy Major, Helmut J Koester, and Yitzhak Schiller. Nmda spikes in basal dendrites of cortical pyramidal neurons. Nature, 404(6775):285–289, 2000.

Erin M Schuman and Daniel V Madison. Locally distributed synaptic potentiation in the hippocampus. Science, 263(5146):532–536, 1994.

Harel Z Shouval, Mark F Bear, and Leon N Cooper. A unified model of nmda receptor-dependent bidirectional synaptic plasticity. Proceedings of the National Academy of Sciences, 99(16):10831–10836, 2002.

Alcino J Silva, Richard Paylor, Jeanne M Wehner, and Susumu Tonegawa. Impaired spatial learning in α-calcium-calmodulin kinase ii mutant mice. Science, 257(5067):206–211, 1992.

Calum S Skene, Maximilian F Eggl, and Peter J Schmid. A parallel-in-time approach for accelerating direct-adjoint studies. Journal of Computational Physics, 429:110033, 2021.

KE Sorra and Kristen M Harris. Occurrence and three-dimensional structure of multiple synapses between individual radiatum axons and their target pyramidal cells in hippocampal area ca1. Journal of Neuroscience, 13(9):3736–3748, 1993.

Luc Stoppini, P-A Buchs, and Dominique Muller. A simple method for organotypic cultures of nervous tissue. Journal of neuroscience methods, 37(2):173–182, 1991.

Soon-Eng Tan and Keng-Chen Liang. Spatial learning alters hippocampal calcium/calmodulin-dependent protein kinase ii activity in rats. Brain research, 711(1-2):234–240, 1996.

Rudi Tong, Thomas Edward Chater, Nigel John Emptage, and Yukiko Goda. Heterosynaptic cross-talk of pre-and postsynaptic strengths along segments of dendrites. Cell Reports, 34(4):108693, 2021.

Jochen Triesch, Anh Duong Vo, and Anne-Sophie Hafner. Competition for synaptic building blocks shapes synaptic plasticity. Elife, 7:e37836, 2018.

Hidetoshi Urakubo, Miharu Sato, Shin Ishii, and Shinya Kuroda. In Vitro Reconstitution of a CaMKII Memory Switch by an NMDA Receptor-Derived Peptide. Biophysical Journal, 106(6):1414–1420, March 2014. ISSN 0006-3495. doi: 10.1016/j.bpj.2014.01.026.

Gayle M Wittenberg and Samuel S-H Wang. Malleability of spike-timing-dependent plasticity at the ca3–ca1 synapse. Journal of Neuroscience, 26(24):6610–6617, 2006.

Sho Yagishita, Akiko Hayashi-Takagi, Graham CR Ellis-Davies, Hidetoshi Urakubo, Shin Ishii, and Haruo Kasai. A critical time window for dopamine actions on the structural plasticity of dendritic spines. Science, 345(6204):1616–1620, 2014.

Shao-Nian Yang, Yun-Gui Tang, and Robert S Zucker. Selective induction of ltp and ltd by postsynaptic [ca2+] i elevation. Journal of neurophysiology, 81(2):781–787, 1999.

Ryohei Yasuda and Hideji Murakoshi. The mechanisms underlying the spatial spreading of signaling activity. Current opinion in neurobiology, 21(2):313–321, 2011.

Lorric Ziegler, Friedemann Zenke, David B Kastner, and Wulfram Gerstner. Synaptic consolidation: from synapses to behavioral modeling. Journal of Neuroscience, 35(3):1319–1334, 2015.

